# Ambra1 haploinsufficiency results in metabolic alterations and exacerbates age-associated retinal degeneration

**DOI:** 10.1101/2022.02.04.476630

**Authors:** Ignacio Ramírez-Pardo, Beatriz Villarejo-Zori, Juan Ignacio Jiménez-Loygorri, Elena Sierra-Filardi, Sandra Alonso-Gil, Guillermo Mariño, Pedro de la Villa, Patrick S Fitze, José Manuel Fuentes, Ramón García-Escudero, Raquel Gomez-Sintes, Patricia Boya

**Author notes:** Corresponding authors; Patricia Boya and Raquel Gomez-Sintes. Shared first authorship (Ignacio Ramírez-Pardo, Beatriz Villarejo-Zori).

## Abstract

Autophagy is a key process in the maintenance of cellular homeostasis. The age-dependent decline in retinal autophagy has been associated with photoreceptor degeneration. Retinal dysfunction can also result from damage to the retinal pigment epithelium (RPE), as the RPE–retina constitutes an important metabolic ecosystem that must be finely tuned to preserve visual function. While studies of mice lacking essential autophagy genes have revealed a predisposition to retinal degeneration, the consequences of a moderate reduction in autophagy, similar to that which occurs during physiological aging, remain unclear. Here, we describe a retinal phenotype consistent with accelerated aging in mice carrying a haploinsufficiency for *Ambra1*, a pro-autophagic gene. These mice show protein aggregation in the retina and RPE, metabolic underperformance, and premature vision loss. Moreover, *Ambra1^+/gt^* mice are more prone to retinal degeneration after RPE stress. These findings indicate that autophagy provides crucial support to RPE-retinal metabolism and protects the retina against stress and physiological aging.

## Introduction

The eye is a very metabolically active tissue with high levels of nutrient and oxygen consumption. The avascular outer retina relies on nutrient diffusion from the choroid vessels through the retinal pigmented epithelium (RPE), which supplies the retina with nutrients such as glucose [1, 2]. This glucose is then metabolized via the tricarboxylic acid (TCA) cycle, oxidative phosphorylation (OXPHOS), and aerobic glycolysis to produce glycerol and ATP, which in turn support phototransduction and phospholipid synthesis to renew photoreceptor outer segments (POS) [3]. β-oxidation of fatty acids and lactate are the main energy sources of the RPE, and are derived from the digestion of POS and photoreceptor glycolysis, respectively [4]. This arrangement results in a high degree of metabolic interdependence between the neural retina and RPE, whereby glucose is converted into lactate in the retina and then shunted to the RPE [5]. The metabolic ecosystem resulting from this interdependence between the two tissue types is key to the preservation of vision [5]. RPE damage has been implicated in retinal diseases such as age-related macular degeneration (AMD), which leads to impaired visual function and blindness, affects millions globally, and is projected to double in incidence by 2040 [6]. While aging is the primary risk factor for this disease, other processes such as oxidative damage and inflammation have also been implicated in AMD [7].

Autophagy is crucial to preserve cell homeostasis, serves as a recycling mechanism to extract nutrients and eliminate damaged cell components, and has important implications in neurological diseases [8, 9]. Studies are beginning to unravel the role of autophagy in the eye and the consequences of autophagy alterations in ocular and retinal diseases [10, 11]. We previously demonstrated a progressive, age-dependent decrease in autophagic flux in the retina of C57/BL6 mice, beginning at 12 months of age [12]. In addition, complete autophagy deficiency, achieved by deleting Atg5 in neuronal precursors, resulted in severe retinal degeneration and vision loss at 7 weeks of age [12] while Atg5 deletion in only cones or rods resulted in much more subtle phenotypes [10,13,14]. Two mouse models that completely lack the essential autophagy proteins Atg5 or Atg7 in the RPE show increased predisposition to age-related retinal degeneration [15]. However, other studies in which these same autophagy regulators were eliminated in the mouse RPE reported no retinal degeneration [16, 17], indicating that further research is needed to determine whether complete autophagy blockade in the RPE results in reduced visual function. More importantly, it is unclear how slight reductions in autophagy in both the RPE and retina, as occurs during physiological aging, affect retinal metabolism and visual function.

AMBRA1 (activating-molecule in Beclin-1-regulated autophagy) is a multifunctional scaffold protein that promotes autophagy initiation by triggering BECLIN1 and VPS34 interaction [18, 19] and ULK1 activity in a mTOR-dependent manner [20]. *Ambra1*-deficient (*Ambra1^gt/gt^*) mice display embryonic lethality and show massive alterations in the central nervous system (CNS) including exencephaly and spina bifida [18], as well as neurogenesis alterations in the olfactory bulb and sub-ventricular zone [21, 22]. Moreover, decreased *Ambra1* expression is linked to an attenuated autophagic response that may be implicated in neural disorders and neurodegenerative diseases such as autism, schizophrenia, and Alzheimeŕs disease [23–25]. Recent findings indicate that AMBRA1 mediates an autophagy-mediated protective response to injury in retinal ganglion cells (RGC) [26, 27]. It remains to be determined how *Ambra1* deficiency affects the retina-RPE ecosystem in the context of aging.

Here, we show that monoallelic deletion of *Ambra1* results in diminished autophagic flux and induces protein aggregation, lipid peroxidation, and oxidative stress in the retina and RPE. Loss of autophagy-dependent proteostasis leads to exacerbated RPE degeneration at 1 year of age and retinal neurodegeneration in 2-year-old mice, culminating in premature vision loss. Moreover, autophagy reduction is associated with metabolic imbalance in *Ambra1^+/gt^* mice, including mitochondrial defects, altered glycolysis, and energetic underperformance. These data demonstrate for the first time that partial loss of autophagy results in marked metabolic alterations and increased susceptibility to AMD-like RPE stress and age-dependent retinal degeneration.

## Results

### Age-dependent vision loss is exacerbated in Ambra1^+/gt^ mice

We investigated how a moderate reduction in autophagy, as occurs during physiological aging, impacts retinal homeostasis. We used *Ambra1^+/gt^* haploinsufficient mice [18], which are viable and display slight reductions in autophagic flux [27]. This outbred model, which has a CD1 background, provides a suitable means of studying age-dependent changes in a physiologically relevant context [28]. Autophagic flux decreased with age in this mouse strain, in agreement with our previous findings (Fig. 1A and S1B) [12]. Retinas from *Ambra1^+/gt^* mice displayed reduced *Ambra1* gene expression (Fig. S1A) and a parallel reduction in autophagic flux compared with control (*Ambra1^+/+^*) littermates at a young age (3–4 months old) (Fig. 1A). Compared with control littermates, gene expression of many essential autophagy regulators such as *Atg5* and *Sqstm1* was already diminished in middle-aged (12-15 months) *Ambra1^+/gt^* mice, and further decreased in old (22–26 months) mice (Fig. 1B). At the protein level, Atg5-12 conjugation or unconjugated-Atg5 were decreased slightly in parallel with the age-associated decline in autophagy. *Ambra1* haploinsufficiency was not associated with any alterations in Atg5 forms (Fig. S1C). Moreover, *Ambra1^+/gt^* mice displayed a sustained reduction in mRNA expression of genes such as *Tfeb* and *Wipi2* (Fig. 1B), suggesting exacerbation of autophagy deficiency in old *Ambra1^+/gt^* mice. Histological analysis revealed a progressive reduction of LC3^+^ puncta with increasing age in all retinal layers, an effect that was more pronounced in *Ambra1^+/gt^* mice (Fig. 1C, E and S1E). In agreement with the observed reduction in autophagic flux in young animals, *Ambra1^+/gt^* mice showed a greater accumulation of protein aggregates as measured using the ProteoStat® protein aggregation assay (Fig. 1D, F). These differences were abolished with increasing age: in control *Ambra1^+/+^* littermates protein aggregation increased with age. Together, these results demonstrate an age-associated decline in autophagy function that correlates with the loss of proteostasis, a phenotype already evident in young *Ambra1^+/gt^* mice.

**Figure 1.**
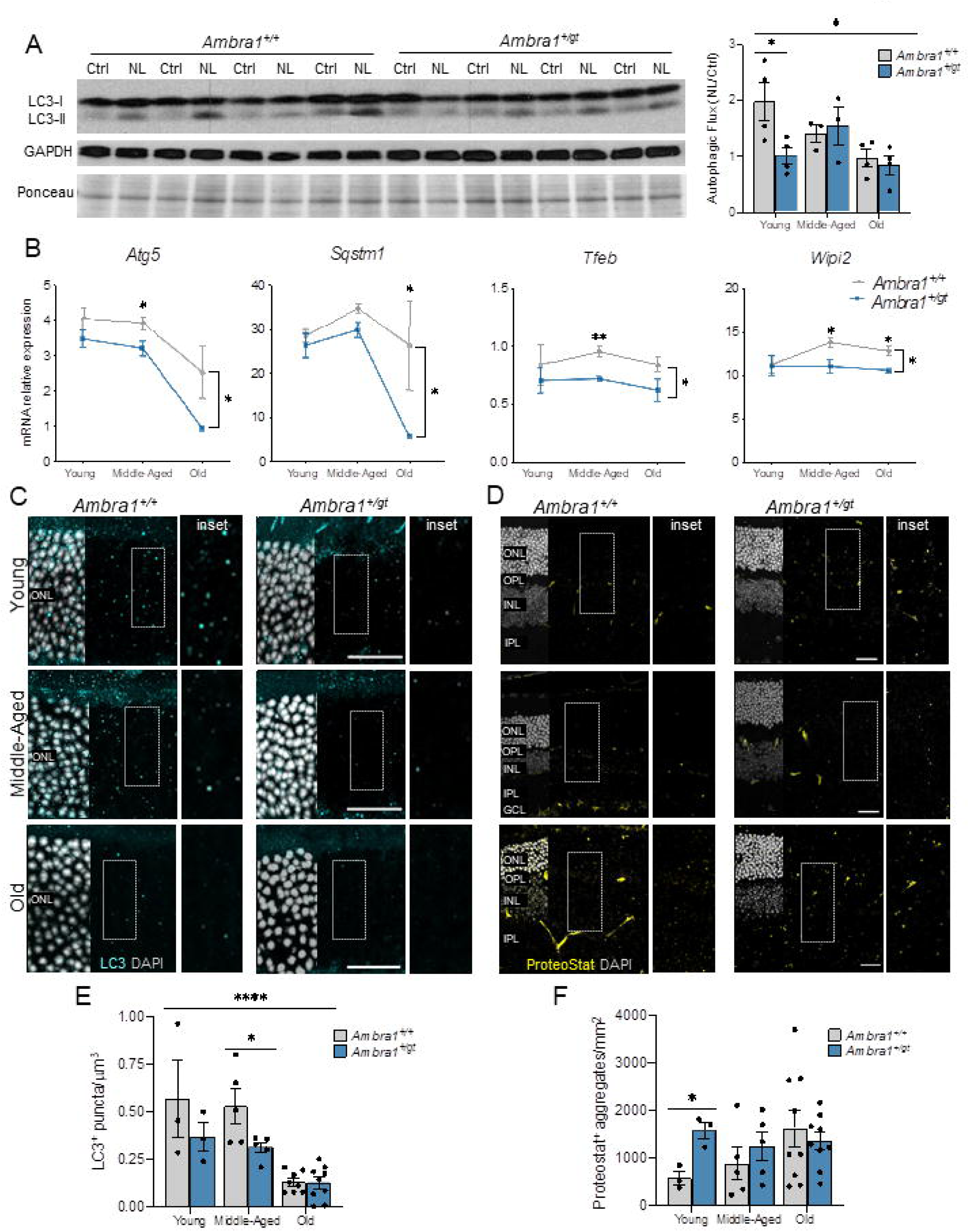
*Ambra1^+/gt^* retinas show a pronounced decrease in autophagy and increased protein aggregation. (**A**) Changes in autophagic flux determined by Western blot for LC3-II in the absence or presence of NH_4_Cl and leupeptin (NL, to block lysosomal degradation) in whole retina extracts from young *Ambra1*^+/+^ and *Ambra1^+/gt^* mice. Quantification of autophagic flux in young (3 months), middle-aged (MA) (12-15 months), and old mice (22-26 months) is shown in the right-hand panel. Western blots corresponding to middle-aged and old mice are shown in Fig. S1. Autophagic flux is determined as the ratio between NL and Control (n = 3–4 per group). (**B**) Decreased mRNA expression of *Atg5*, *Sqstm1*, *Tfeb* and *Wipi2* as determined by qPCR of whole retina extracts from young, middle-aged, and old *Ambra1*^+/+^ and *Ambra1*^+/gt^ mice (n = 3–4 per group). (**C-F**) Representative images and quantification of LC3 puncta in the outer nuclear layer (cyan) (**C**) and protein aggregates in the whole retina as measured by ProteoStat® protein aggregation assay (yellow) (**D**) in young, middle-aged, and old *Ambra1*^+/+^ and *Ambra1^+/gt^* mice. Nuclei were counterstained with DAPI (grey). Graphs show quantification of LC3^+^ puncta (**E**) and protein aggregates (**F**) (n = 3–9 per group). Data are presented as the mean ± SEM. *p <0.05: two-tailed Student’s *t*-test or Mann-Whitney *U*-test (A, E, F); two-way ANOVA followed by *post hoc* LSD test for age (A, E) and genotype (**B**). Scale bars: 25 µm.

Defective autophagy in the *Ambra1^+/gt^* retina was accompanied by reduced visual function, an effect that was already significant in middle-aged animals compared with control littermates (Fig. 2A-H). In low light (scotopic) conditions, we observed decreased amplitudes of b-scot (low intensity stimulus) and both a- and b-mixed (high intensity stimulus) waves, implying decreased photoreceptor and bipolar cell signal transduction, respectively, in *Ambra1^+/gt^* versus *Ambra1^+/+^* mice (Fig. 2A and B). In photopic conditions, decreased amplitudes of b-photopic and flicker waves indicated defective cone-dependent responses (Fig. 2F and G). More detailed analysis based on b:a wave ratio showed a significant reduction in old *Ambra1^+/gt^* versus *Ambra1^+/+^* animals (Fig. 2D), suggesting poor performance of bipolar cells at this age due to a more pronounced reduction of b-wave than a-wave amplitudes. Finally, diminished oscillatory potentials (OP) suggested poor performance of inner layer cells (Fig. 2E). Regression analyses revealed that monoallelic deletion of *Ambra1* resulted in faster age-associated vision loss than in control mice, as evidenced by a significantly steeper slope for *Ambra1^+/gt^* versus *Ambra1^+/+^* mice for all parameters studied (Fig. S2).

**Figure 2.**
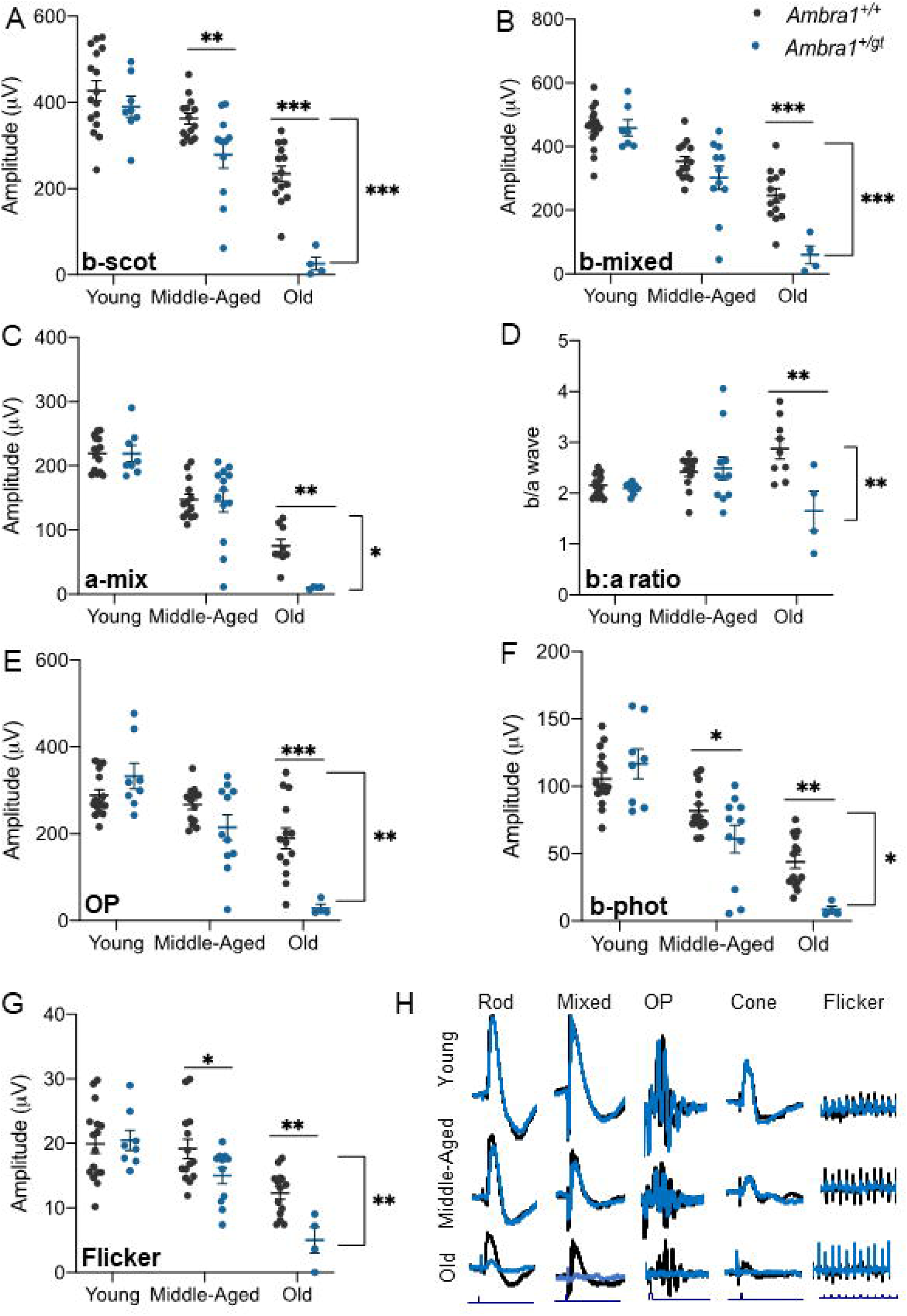
Age-dependent vision loss is exacerbated in *Ambra1^+/gt^* mice. Visual function assessed by measuring electroretinographic responses in young, middle-aged, and old *Ambra1*^+/+^ and *Ambra1^+/gt^* mice. (**A**) b-scotopic (b-scot), (**B**) b-mixed (b-mix), and (**C**) a-mixed (a-mix) waves, and (**D**) b:a wave ratio and (**E**) oscillatory potentials (OP) were measured in dark-adapted conditions, and (**F**) b-photopic (b-phot) and (**G**) flicker waves were measured in light-adapted conditions. (**H**) Representative electroretinographic recording waves of young (n=11–16), middle-aged (n=8–14), and old (n=4–14) *Ambra1*^+/+^ and *Ambra1^+/gt^* mice. Data are presented as the mean ± SEM. *p <0.05, **p <0.01, ***p <0.001: two-way ANOVA (bracket bars) followed by Fischer’s LSD *post hoc* test for genotype (**A–H**).

To examine the effects of the age-associated decline in autophagy on retinal morphology, we compared ONL (outer nuclear layer) nuclear density in young, middle-aged, and old *Ambra1^+/gt^* mice versus *Ambra1^+/+^* littermates. Old *Ambra1^+/gt^* mice showed reduced ONL volume density (Fig. 3A,B), but no significant increase in the number of apoptotic TUNEL-positive cells (Fig. S3A). These apoptotic cells may have been already removed by the more numerous microglial cells with amoeboid morphology, often associated to their phagocytic phenotype (Fig. S3C). The number of cones stained with either cone arrestin or red/green opsin was reduced in aged *Ambra1^+/gt^* retinas (Fig. 3C,D and S3B), indicating that *Ambra1* haploinsufficiency results in an age-associated reduction in cone population. We next investigated whether monoallelic loss of *Ambra1* results in alterations in the inner retina. *Ambra1^+/gt^* mice showed a reduction in the thickness of the inner plexiform layer (IPL) (Fig. 3E, F) compared with *Ambra1^+/+^* littermates, suggesting reduced synapsis between retinal neurons. Moreover, in these mice, bipolar cells showed protrusions towards the ONL (Fig. 3E [insets], G), a phenotype associated with the bipolar cell response to photoreceptor retraction and connectivity loss [29]. In terms of photoreceptor and bipolar morphology and connectivity, the observed phenotypes match perfectly with the loss visual function revealed by electroretinogram. Old *Ambra1^+/gt^* and control *Ambra1^+/+^* mice showed no significant differences in the number of retinal ganglion cells stained with the specific transcription factor POU4F1, also known as BRN3A (Fig. 3H).

**Figure 3.**
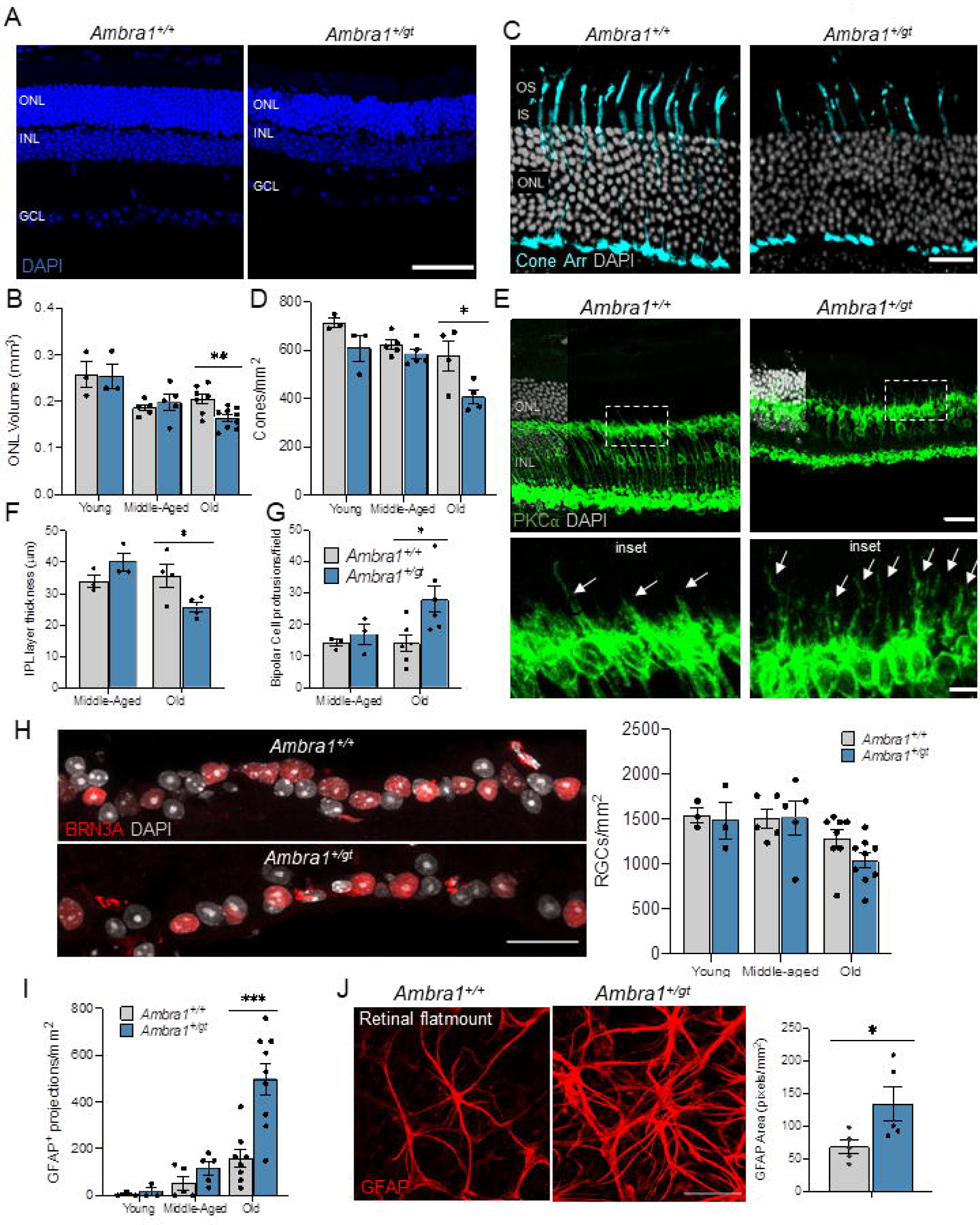
Retinal degeneration is observed in old *Ambra1^+/gt^* retinas. (**A**) Representative images of retinal nuclei (DAPI, blue). (**B**) Graph depicts changes in ONL volume in *Ambra1^+/+^* and *Ambra1^+/gt^* (n = 3–8 per group) mice over time. (**C**) Cone morphology evaluated using cone arrestin immunostaining (cyan) and (**D**) corresponding quantification in young, middle-aged, and old *Ambra1^+/+^* and *Ambra1^+/gt^* mice (n = 3– 5 per group). (**E**) Immunostaining of bipolar cells (protein kinase C [PKC], green) in old *Ambra1^+/+^* and *Ambra1^gt/+^* mice. Insets (below) show protrusions from bipolar cells towards the outer nuclear layer (ONL) (arrows). (**F**) Quantification of inner plexiform layer (IPL) thickness (n = 3–4 per group). (**G**) Determination of number of protrusions from bipolar cells (n = 3–4 per group). (**H**) Immunostaining of retinal ganglion cells with Brn3a (red) and corresponding quantification in young, middle-aged, and old *Ambra1*^+/+^ and *Ambra1*^+/gt^ retinas (n = 3–8). (**I**) Quantification of GFAP-positive projections towards the outer retina in each experimental group (n = 3–9). (**J**) GFAP immunostaining (red) of retinal flat mounts from old *Ambra1^+/+^* and *Ambra1^+/gt^* mice and quantification of GFAP-positive area (n = 5 per group). Data are presented as the mean ± SEM. *p <0.05, **p <0.01, ***p <0.001: two-tailed Student’s *t*-test (**B, D, F, G, H, I, J**). Scale bars: 50 µm (**A, C**); 25 µm (**C, H, J**); 10 µm (insets in **E**).

Müller cells are retinal glial cells that provide trophic factors and support the neuroretina. In response to damage stimuli, Müller cells activate and extend GFAP-positive projections towards the outer retina. A persistent inflammatory and gliotic response accompanying aging has been described in mice [30]. Although we did not observe major morphological changes in Müller glia stained with the Müller cell marker glutamine synthetase (GS) (Fig. S3D), we observed a massive increase in the number of GFAP projections towards the ONL in aged *Ambra1^+/gt^* versus *Ambra1^+/+^* mice (Fig. 3I). This phenotype was accompanied by a larger area of GFAP^+^ staining at the level of the ganglion cell layer in old *Ambra1^+/gt^* retinal flat mounts (Fig. 3J), indicating astrocyte hypertrophy. In agreement with this gliosis response, we observed a slight increase in amoeboid microglia in old *Ambra1^+/gt^* versus *Ambra1^+/+^* retinas (Fig. S3C) that could be linked to their activated state during inflammation [31, 32]. Taken together, these data suggest that *Ambra1* haploinsufficiency results in an accelerated retinal aging phenotype, accompanied by a proinflammatory state and vision defects.

### Monoallelic deletion of Ambra1 results in metabolic alterations in the retina associated with mitochondrial dysfunction

Recent findings by our group demonstrate that autophagy-deficient animals display marked metabolic alterations in the embryonic retina [33], and that retinal degeneration is often associated with metabolic imbalances [34]. We next explored how deficient autophagy influences the retinal metabolome in the early stages of degeneration by comparing retinas from young and middle-aged *Ambra1^+/+^* and *Ambra1^+/gt^* mice. Metabolite levels were determined by liquid chromatography/gas chromatography– mass spectrometry (LC/GC–MS) and concentrations represented as the area under the curve and metabolite ratios. Although sample correlation, principal component analysis, and unsupervised hierarchical clustering analysis revealed greater differences for age than for genotype (Fig. 4A, and S4A-B), middle-aged *Ambra1^+/gt^* retinas showed the most pronounced changes in retinal metabolome.

**Figure 4.**
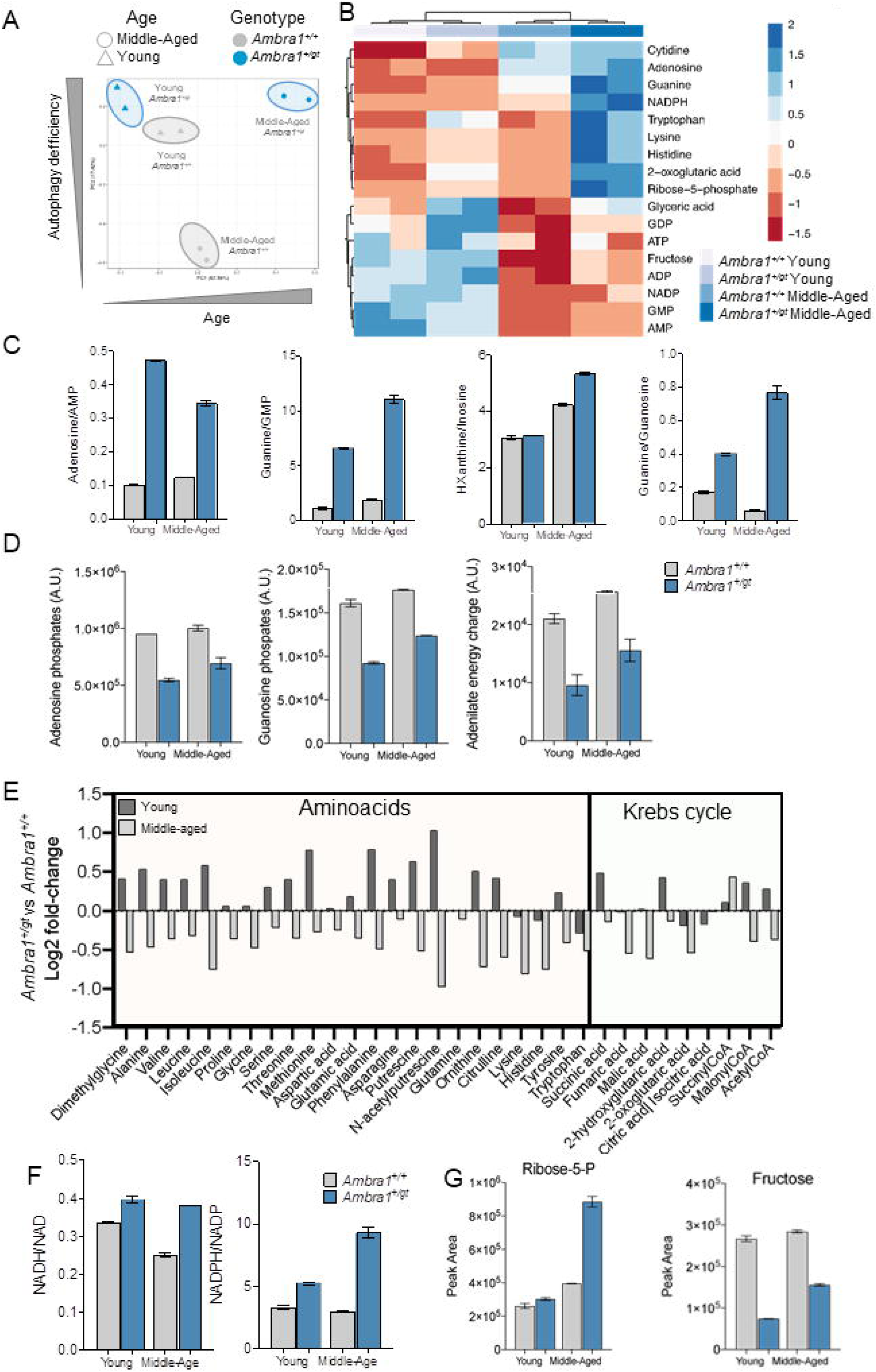
Metabolic imbalance in *Ambra1^+/gt^* mice. (**A**) Principal component analysis (PCA) of retinal mentabolites in samples from young and middle-aged *Ambra1^+/+^* and *Ambra1^+/gt^* retinal metabolites. (**B**) Heatmap and hierarchical clustering (HCL) analysis of metabolites for which statistically significant alterations were observed in *Ambra1^+/gt^* mice (n = 2 pools of 4 retinas per group). Statistical significance (Skillings-Mack test) was set at p <0.05. (**C**) Purine metabolism metabolites as determined by the ratios of related metabolic compounds in young and middle-aged *Ambra1^+/+^* and *Ambra1^+/gt^* retinas (n = 2 per group). (**D**) Pools of charged purine (adenosine and guanosine) phosphate nucleotides and estimation of the adenylate energy charge in young and middle-aged *Ambra1^+/+^* and *Ambra1^+/gt^* retinas (n = 2 per group). (**E**) Fold changes in the levels of amino acids and Krebs cycle-related metabolites in young and middle-aged *Ambra1^+/+^* versus *Ambra1^+/gt^* mice (n = 2 per group). (**F**) Retinal redox status as measured by the ratios of NAD-related metabolites in young and middle-aged *Ambra1^+/+^* versus *Ambra1^+/gt^* littermates (n = 2 per group). (**G**) Changes in the pentose-phosphate pathway as determined based on relative levels of representative metabolites (fructose and ribose-5-phosphate) in young and middle-aged *Ambra1^+/+^* versus *Ambra1^+/gt^* littermates (n = 2 per group). Data are presented as the mean ± SEM.

To determine which metabolites were dysregulated, we performed a Skillings-Mack analysis to identify significant genotype-related effects, followed by an enrichment analysis. Of the 100 metabolites measured (Fig. S4A), 16 were significantly altered in middle-aged *Ambra1^+/gt^* retinas (Fig. 4B). Those metabolites are implicated in purine metabolism, the pentose phosphate pathway, and the Warburg effect, among other processes (Fig. S4C). Nucleotide metabolite ratios revealed alterations in adenine and guanine metabolism in *Ambra1^+/gt^* versus *Ambra1^+/+^* retinas (Fig. 4C).

Furthermore, middle-aged *Ambra1^+/gt^* retinas displayed marked reductions in total nucleotide phosphate levels and adenylate energy charge (Fig. 4D), a phenotype associated with energetic underperformance [35]. Interestingly, while amino acid levels were increased in young *Ambra1^+/gt^* versus *Ambra1*^+/+^ mice, levels of most amino acids analyzed and of many TCA cycle intermediates were reduced in middle-aged *Ambra1^+/gt^* mice (Fig. 4E). Finally, the NADH:NAD and the NADPH:NADP ratios were altered and levels of ribose-5-phosphate increased in middle-aged *Ambra1^+/gt^* versus *Ambra1*^+/+^ mice (Fig. 4F-G), indicating marked metabolic alterations suggestive of defective oxidative metabolism in *Ambra1^+/gt^* mice.

We next investigated whether alterations in mitochondrial function in *Ambra1*-deficient retinas could explain the exacerbated metabolic dysregulation observed in middle-aged animals. Surprisingly, marked reductions in mitochondrial membrane potential as determined by flow cytometry with DiOC6 were already detectable in young *Ambra1^+/gt^* retinas (Fig. 5A). TOMM20 immunostaining revealed augmented mitochondrial mass in most retinal cell layers in aged *Ambra1^+/gt^* animals (Fig. 5B, C). Increases in mitochondrial biogenesis can often compensate for mitochondrial alterations. However, *Ambra1^+/gt^* animals displayed no such changes and old animals even showed reduced mRNA expression of the main mitochondrial biogenesis regulators *Ppargc1a, Nrf1*, *Nfe2l2,* and *Tfam* (Fig. 5D), as well as decreased expression of mitochondrial genes such as *Timm23* and *COXIV* (Fig. 5E). Thus, the increased mitochondrial mass in the absence of biogenesis suggests a defect in the removal of dysfunctional mitochondria via mitophagy in aged *Ambra1^+/gt^* animals.

**Figure 5.**
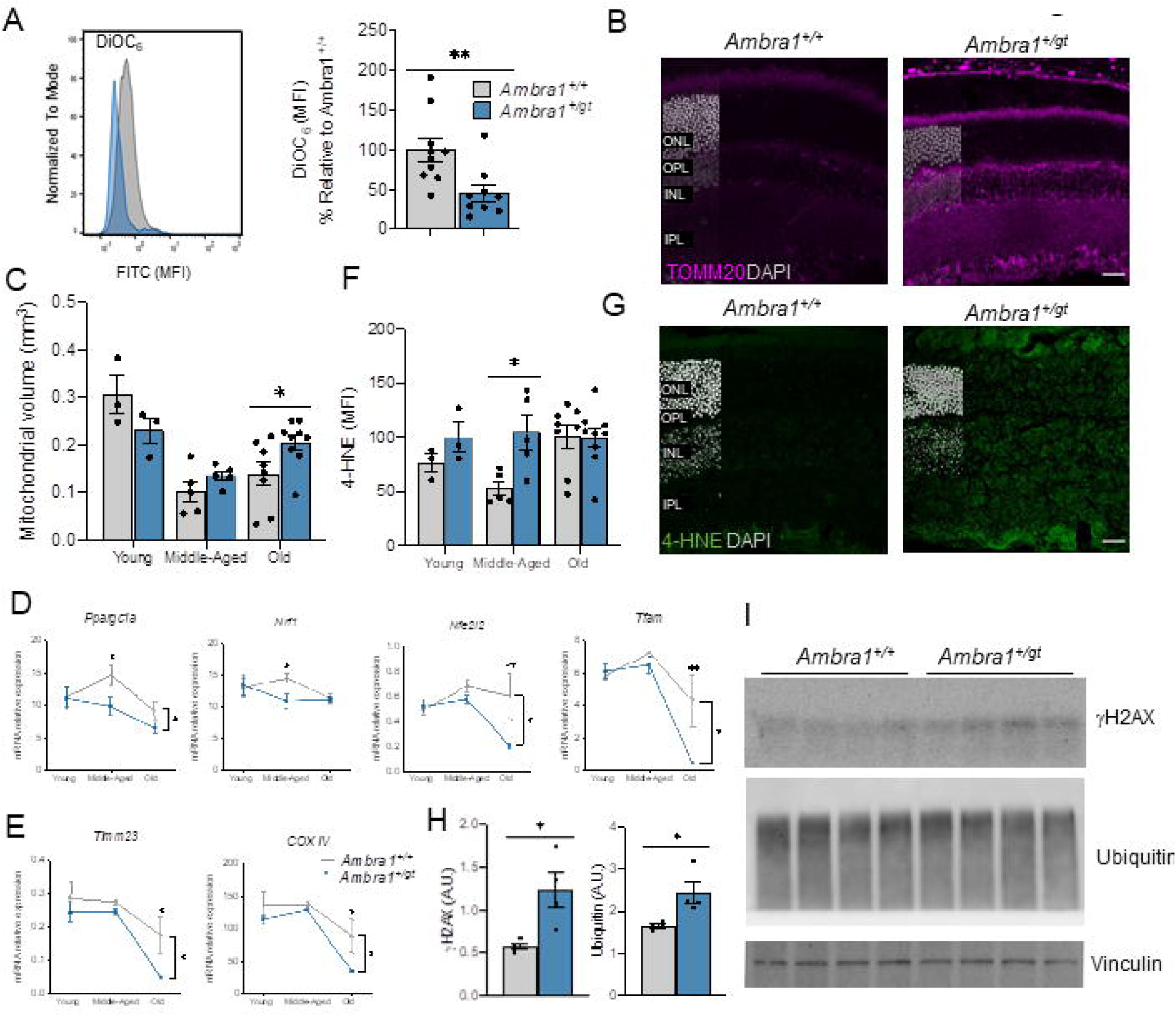
Mitochondrial alterations, cellular damage, and oxidative stress in *Ambra1^+/gt^* retinas. (**A**) Fluorescence-activated cell sorting (FACS) histogram (left) and quantification (right) of mitochondrial membrane potential measured by DiOC6(3) probe in dissociated retinas from young *Ambra1^+/+^* and *Ambra1^+/gt^* mice (n = 9–10 per group). (**B–C**) Immunostaining of mitochondrial mass (TOMM20, magenta). (**B**) Representative image of old *Ambra1^+/+^* and *Ambra1^+/gt^* retinas and (**C**) quantification of mitochondrial volume in young, middle-aged, and old *Ambra1^+/+^* and *Ambra1^+/gt^* retinas (n = 3–8 per group). (**D–E**) Mitochondrial transcriptional signature is decreased in *Ambra1^+/gt^* retinas. Graphs depict the transcriptional levels of mitochondrial biogenesis related-genes (*Ppargc1a*, *Nrf1*, *Nfe2l2*, and *Tfam*) (**D**) and mitochondrial mass-related genes (*Timm23* and *CoxIV*) (**E**). (**F-G**) Immunostaining to determine lipid peroxidation levels (4-HNE, green) in middle-aged and *Ambra1^+/+^* and *Ambra1^+/gt^* retinas and subsequent quantification of 4-HNE mean fluorescence intensity (MFI) in the ONL of young, middle-aged, and old *Ambra1^+/+^* and *Ambra1^+/gt^* littermates (n = 3– 8 per group). Nuclei are counterstained with DAPI (grey). (**H-I**) Protein levels of γ-H2AX, ubiquitin, and vinculin (loading control) in middle-aged *Ambra1^+/+^* and *Ambra1^+/gt^* retinas as determined by Western blot and quantification of protein levels is shown on the right (n = 4 per genotype). Data are presented as the mean ± SEM. *p <0.05, **p <0.01: two-tailed Student’s *t*-test or Mann-Whitney *U*-test (**A**, **C**, **F**, **H**) and two-way ANOVA followed by Fisher’s LSD *post hoc* test for genotype (**D, E**). Scale bars: 25 µm.

We next investigated whether these mitochondrial alterations result in oxidative damage to intracellular membranes. Photoreceptor outer segments are enriched in lipids that are susceptible to peroxidation upon oxidative stress, affecting their functionality [36]. Compared with control littermates, middle-aged *Ambra1^+/gt^* mice showed increased levels of lipid peroxidation in all retinal layers, as determined by measuring 4-hydroxynonenal (4-HNE) (Fig. 5F, G), and other signs of cellular damage such as increased γH2AX and ubiquitin levels (Fig. 5H, I). Together, these data indicate marked mitochondrial alterations in *Ambra1*-deficient retinas, including decreased mitochondrial membrane potential in young animals that cannot be compensated for in later life by autophagy or mitochondrial biogenesis, potentially resulting in lipid peroxidation and oxidative damage to cellular membranes.

Our enrichment analysis (Fig. S4C) revealed upregulation of the pathway underlying the Warburg effect. Because the pyruvate:lactate ratio was increased in young *Ambra1^+/gt^* versus *Ambra1*^+/+^ mice (Fig. 6A), we next examined mRNA expression of the main glycolytic enzymes. In all age groups, *Ambra1^+/gt^* retinas showed reduced mRNA expression of many glycolytic enzymes, including *Gapdh, Hk2,* and *Pkm* (Fig. 6B), and a tendency towards reduced GAPDH and HK2 immunofluorescence (Fig. 6C-F). In agreement with our metabolic findings in middle-aged *Ambra1^+/gt^* retinas, we observed decreased transcription of enzymes that help maintain cellular redox status. For example, we observed reduced expression of the NAD(P)H quinone dehydrogenase 1 *(Nqo)1* (Fig. 6G), an enzyme that oxidizes NADH to NAD in the electron transport chain, in agreement with the aforementioned alterations in NADH:NAD ratio (Fig. 4F). Furthermore, we detected lower mRNA levels of the pyruvate dehydrogenase (E1) alpha subunit gene (*Pdha1)*. PDHA1 protein is crucial to allow entry of acetyl-CoA produced from pyruvate into the TCA cycle, where it serves as a source of carbon for anabolism (Fig. 6G). Together these data demonstrate that mitochondrial alterations already evident in young *Ambra1^+/gt^* retinas may underlie the metabolic changes, including decreased glycolysis and energetic underperformance, observed in later life.

**Figure 6.**
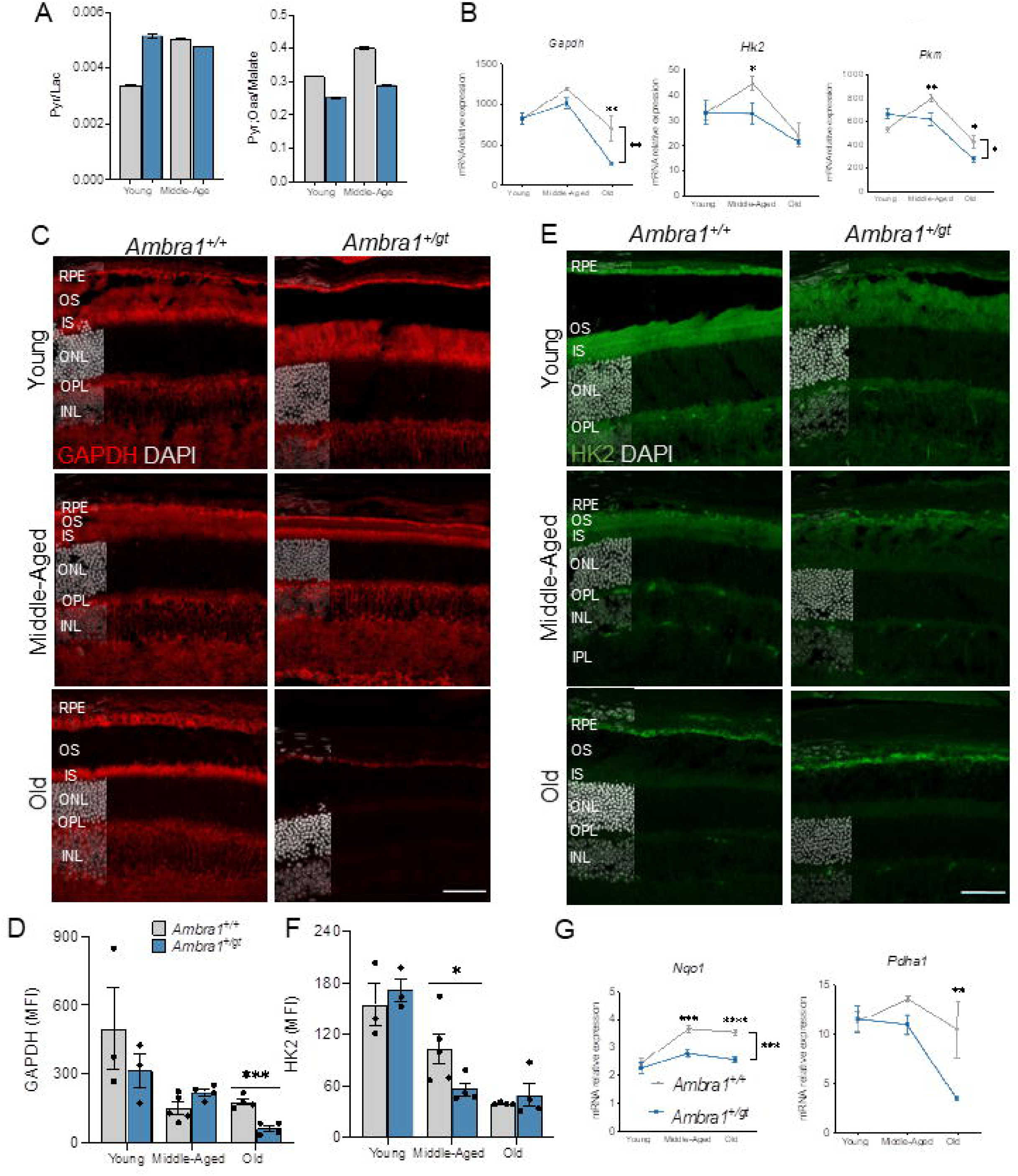
Glycolytic disfunction in *Ambra1^+/gt^* retinas. (**A**) Retinal metabolic status as determined by ratios of glycolytic-end metabolites in young and middle-aged *Ambra1^+/+^* and *Ambra1^+/gt^* mice (n = 2 per group). **(B)** mRNA expression of glycolysis-related genes (*Gapdh, Hk2*, and *Pkm)* in young, middle-aged, and old *Ambra1^+/+^* and *Ambra1^+/gt^* retinas. (**C**) Immunostaining for GAPDH (red) in young, middle-aged, and old *Ambra1^+/+^* and *Ambra1^+/gt^* retinas (n = 3–4 per group) and (**D**) corresponding quantification. (**E**) Immunostaining for HK2 (green) in young, middle-aged, and old *Ambra1^+/+^* and *Ambra1^+/gt^* retinas (n = 3–4 per group) and (**F**) corresponding quantification. Nuclei in **C** and **E** are counterstained with DAPI (grey). (**G**) mRNA expression of redox status-related enzymes (*Nqo1 and Pdha1)* as determined by qPCR of whole retina extracts from young, middle-aged, and old *Ambra1^+/+^* and *Ambra1^+/gt^* mice (n = 3–4 per group). Data are presented as the mean ± SEM. *p <0.05, **p <0.01, ***p <0.001, ****p <0.0001: two-tailed Student’s *t*-test (**D**, **F**) and two-way ANOVA (bracket bars) followed by Fisher’s LSD *post hoc* test for genotype (**B, G**). Scale bars in **E**, **F**: 50 µm

### RPE alterations in Ambra1^+/gt^ mice result in increased susceptibility to stress

Retinal metabolism relies heavily on the RPE, and vice versa. For example, photoreceptors mainly obtain energy through glycolysis from glucose provided by the RPE, and, in exchange, return lactate and lipid-containing outer segments that are used to fuel oxidative phosphorylation in the RPE [37]. Because alterations in RPE homeostasis have important consequences for retinal metabolism and function, we examined age-associated changes in RPE function in *Ambra1^+/gt^* mice. While the RPE in young *Ambra1^+/gt^* mice showed no alterations with respect to control littermates, phalloidin staining in middle-aged *Ambra1^+/gt^* animals revealed important RPE lesions corresponding to monolayer disruption (Fig. 7A) [38]. Moreover, middle-aged *Ambra1^+/gt^* mice displayed fewer (Fig. 7B) and larger (Fig. 7C) RPE cells, phenotypes associated with RPE cell death, whereby the remaining cells expand to cover the area left by dying cells [38]. ProteoStat protein aggregation assay showed that monoallelic loss of *Ambra1* was associated with increased proteotoxicity, an effect that was already significant in young animals (Fig. 7D). Importantly, those differences were no longer observed in old animals, as protein aggregation was also increased in the old control animals. In agreement, autophagosome number determined by LC3 staining was reduced both in flat mounts and RPE cryosections from *Ambra1^+/gt^* versus *Ambra1^+/+^* mice (Fig. S5A-D). The age-associated increase in lipid peroxidation was significantly augmented in the RPE of middle-aged *Ambra1^+/gt^* mice (Fig. 7E). Chronic RPE dysfunction is associated with the appearance of autofluorescent lipofuscin aggregates [36]. Lambda-scan autofluorescence analysis showed specific accumulation of lipofuscin inside swollen lysosomes in old *Ambra1^+/gt^* mice (Fig. 7F). In agreement, lysosomes in *Ambra1^+/gt^* animals were similar in number to those of control littermates, but with greater individual volume (Fig. 7G, H). In conclusion, these data show that monoallelic loss of *Ambra1* is associated with marked alterations in the RPE from a young age, including increased protein aggregation, oxidative stress, lysosomal alterations, and lipofuscin accumulation, resulting in RPE cell death by middle age.

**Figure 7.**
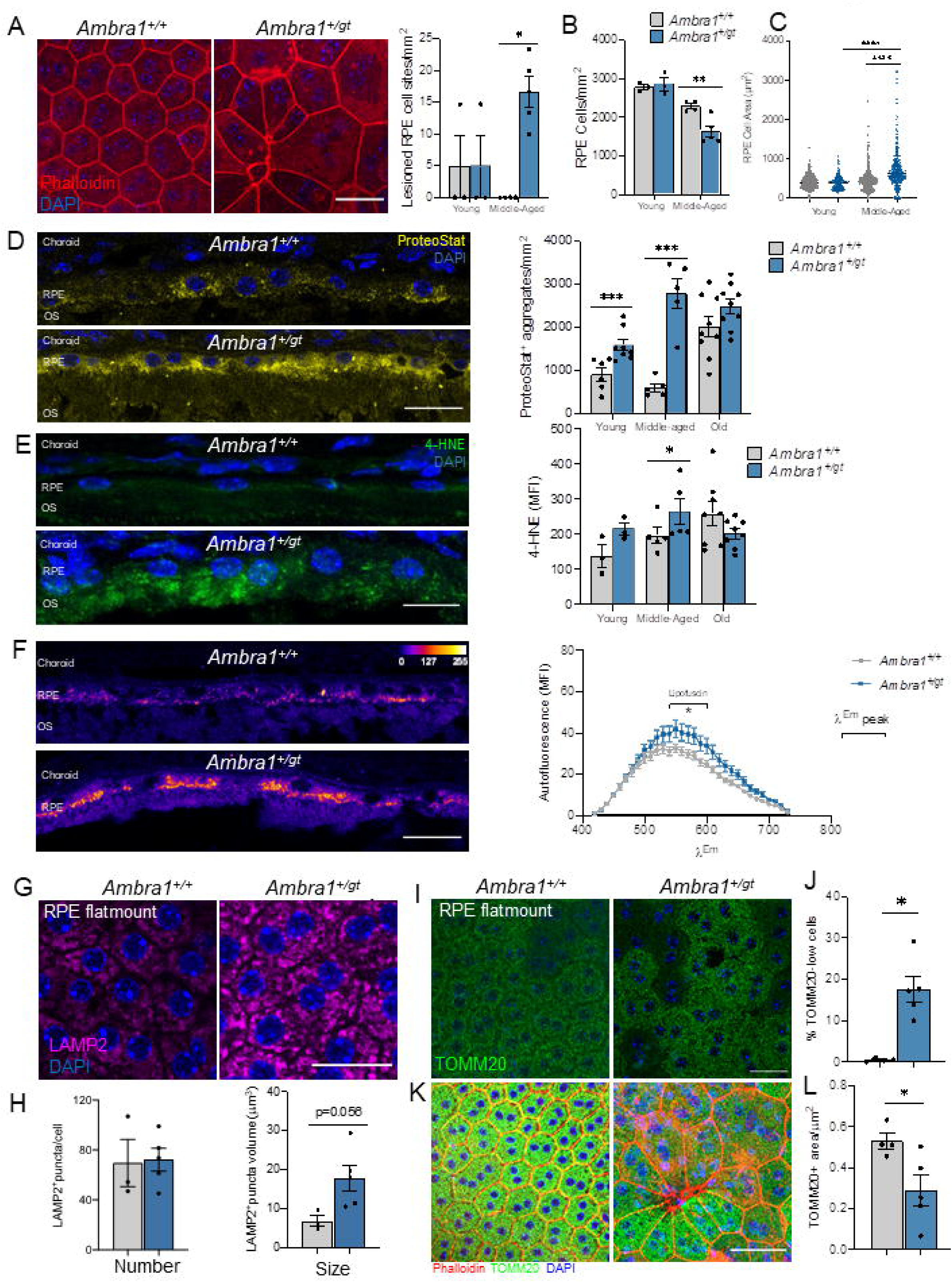
*Ambra1^+/gt^* mice display abnormalities in the retinal pigmented epithelium (RPE) associated with premature aging. (**A**) RPE morphology assessed by phalloidin (red) staining in RPE flat mounts from middle-aged *Ambra1^+/+^* and *Ambra1^+/gt^* littermates, showing increases in cell size and lesion site number in the RPE monolayer in *Ambra1^+/gt^* mice (left). Graph on the right shows the quantification (right) of lesion site number (dying/dead RPE cells; points where >3 cells are in contact) per mm^2^ (n = 3–5 per group). (**B**) Quantification of the number of cells/mm^2^ in RPE flat mounts from young and middle-aged *Ambra1^+/+^* and *Ambra1^+/gt^* littermates (n = 3–5 per group). (**C**) Quantification of the area of individual RPE cells (µm^2^) in RPE flat mounts from young and middle-aged *Ambra1^+/+^* and *Ambra1^+/gt^* littermates (n = 3–5 animals per genotype/age group, >350 cells per group). (**D**) ProteoStat® assay to detect protein aggregation (yellow), showing the accumulation of protein aggregates in the RPE of young, middle-aged, and old *Ambra1^+/+^* and *Ambra1^+/gt^* littermates. Single z-images are shown. Right-hand panel shows quantification of protein aggregates (n = 5–8 per group). (**E**) Lipid peroxidation as determined by 4-HNE staining (green) in young, middle-aged, and old *Ambra1^+/+^* and *Ambra1^+/gt^* littermates and corresponding quantification of mean fluorescence intensity (MFI) (n = 3–5 per group). (**F**) Measurement of autofluorescence in the RPE using Lambda-scan to capture the entire emission spectrum upon UV excitation in old *Ambra1^+/+^* and *Ambra1^+/gt^* littermates (n = 5–8 per group). (**G**) RPE flat mounts from middle-aged *Ambra1^+/+^* and *Ambra1^+/gt^* immunostained with the lysosomal membrane marker LAMP2 (nuclei are counterstained with DAPI [blue]), and (**H**) corresponding quantification of the number and individual volume of puncta (n = 3–5 per group). (**I–K**) Immunostaining of mitochondria (TOMM20, green) in RPE flat mounts from middle-aged *Ambra1^+/+^* and *Ambra1^+/gt^* littermates. Nuclei were counterstained with DAPI (blue) and the F-actin marker phalloidin (red). (**J–L**) Quantification of TOMM20 staining in **I–K**. Data are presented as the mean ± SEM. *p <0.05, **p <0.01; ***p <0.001; ****p <0.0001: two-tailed Student’s *t*-test (**B, D–F**), Mann-Whitney *U*-test (**A, H, J, L**), or Kruskal-Wallis test followed by Dunn’s *post hoc* test (**C**). Scale bars: 25 µm.

As previously stated, mitochondrial function is required to sustain ATP synthesis in the RPE. RPE flat mounts from middle-aged *Ambra1^+/gt^* animals displayed numerous cells with markedly reduced mitochondria mass, as determined by TOMM20 staining (Fig. 7I, J). Those mitochondria-poor cells were enlarged and showed alterations in cytoskeleton architecture, as assessed by phalloidin staining (Fig. 7K, L), suggesting a correlation between reduced mitochondrial mass and RPE hypertrophy. Western blot for TOMM20 and TIMM23 also revealed a trend towards reduced mitochondria protein levels (Fig. S5E). Given the high degree of metabolic interdependence between the retina and RPE, we hypothesize that alterations in mitochondrial function may make the RPE much more dependent on glucose for sustained ATP synthesis. In agreement, our data show a tendency towards increased GLUT1/Slca1 glucose transporter expression in the RPE of *Ambra1^+/gt^* animals, and a tendency towards increased glycolytic GAPDH expression in the RPE of young and middle-aged *Ambra1^+/gt^* mice (Fig. 7F, G), pointing to increased glucose demand in the RPE. These data suggest a scenario in which the autophagy defects in the *Ambra1^+/gt^* RPE result in mitochondrial damage, oxidative stress, and increased glycolysis to fuel ATP synthesis. Finally, the accumulation of activated phagocytic cells was evidenced by an increase in the number of amoeboid IBA1-positive cells in the RPE in middle-aged *Ambra1^+/gt^* mice (Fig. S5H). These data indicate that *Ambra1* haploinsufficiency results in early alterations in proteostasis, lipid peroxidation and reduced mitochondrial mass, leading to RPE dysfunction and death by middle age.

In age-related diseases there is usually an interplay between physiological aging and both genetic and environmental factors. Since the alterations in the RPE of *Ambra1^+/gt^* mice occur earlier and are more severe than those seen in the neuroretina, we used a pharmacological model of RPE degeneration to assess vulnerability to external stressors and age-related diseases in these mice [39, 40]. Middle-aged mice were treated with sodium iodate (SI), a well-studied model of primary RPE damage that leads to secondary retinal degeneration and is often used as a model of dry AMD [39]. Middle-aged *Ambra1^+/gt^* mice and control littermates were intraperitoneally injected with 50 mg/kg SI or vehicle and sacrificed 1 week later. No differences in retinal morphology or retinal layer thickness were observed in the vehicle-treated groups (Fig. 8A-B). However, compared with control littermates, *Ambra1^+/gt^* mice were significantly more sensitive to SI-induced damage, as evidenced by a decrease in ONL thickness (Fig. 8B). Surprisingly, SI-treated *Ambra1^+/gt^* mice also displayed a significant decrease in inner nuclear layer (INL) thickness (Fig. 8B) that was not observed in SI-treated control littermates. This suggests an increased vulnerability of both retinal photoreceptors and interneurons to stress, correlating with loss of visual function in old animals. Supporting increased stress vulnerability in middle-aged *Ambra1^+/gt^* mice, these animals showed increased photoreceptor cell death, as determined by TUNEL staining (Fig. 8C, E), and decreased cone number after SI-induced damage (Fig. 8D, F). SI treatment exerts a degenerative effect following a central to peripheral gradient [39]. Assessment of RPE monolayer morphology using phalloidin staining showed severe RPE disruption, even in the peripheral region of the retina, in SI-treated *Ambra1^+/gt^* mice (Fig. 8G). Given the retinal phenotype observed in *Ambra1^+/gt^* mice, and the fact that SI exerts a time- and region-dependent effect [40], this finding demonstrates increased susceptibility of the *Ambra1^+/gt^* RPE to stress, resulting in exacerbated retinal degeneration.

**Figure 8.**
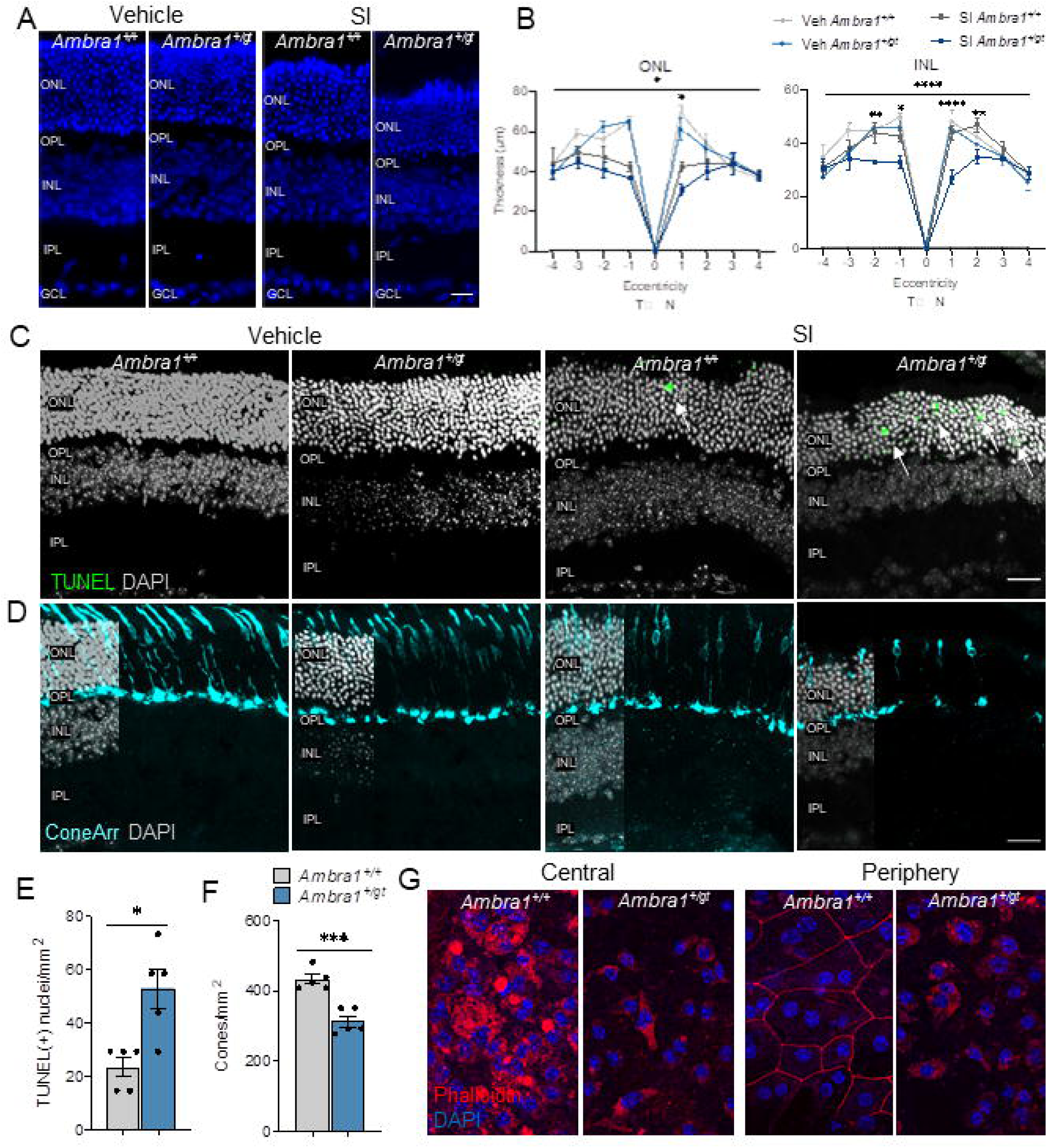
Increased sensitivity to sodium iodate in the retina and RPE of *Ambra1^+/gt^* mice. (**A**) Representative DAPI-stained retinal cryosections from middle-aged *Ambra1^+/+^* and *Ambra1^+/gt^* littermates injected with either 50 mg/kg sodium iodate (SI) or vehicle (PBS) and analyzed after 7 days. (**B**) Quantification of the thickness of the outer nuclear layer (ONL) and inner nuclear layer (INL) of middle-aged *Ambra1^+/+^* and *Ambra1^+/gt^* littermates treated with SI or vehicle (n = 4–5 per group). (**C, D**) Staining of apoptotic cells with TUNEL (**C**, green) and cone arrestin (**D**, cyan) in retinal cryosections from middle-aged *Ambra1^+/+^* and *Ambra1^+/gt^* littermates injected with SI or vehicle. Nuclei were counterstained with DAPI (grey). Single z-images were analyze€(**E**) Quantification of the number of TUNEL-positive photoreceptor nuclei in middle-aged *Ambra1^+/+^* and *Ambra1^+/gt^* littermates treated as in C (n = 5 per group). (**F**) Quantification of the number of cone photoreceptors of middle-aged *Ambra1^+/+^* and *Ambra1^+/gt^* littermates treated as in D (n = 5 per group). (**G**) Phalloidin (red) and DAPI (blue) immunostaining of RPE flat mounts (images from central and peripheral areas are shown) from middle-aged *Ambra1^+/+^* and *Ambra1^+/gt^* littermates injected with 50 mg/kg SI or vehicle. Data are presented as the mean ± SEM. *p <0.05, ***p <0.001, ****p <0.0001: two-way ANOVA followed by Fisher’s LSD *post hoc* test for geno type (**B**); unpaired Student’s *t* test (**D**); or Mann-Whitney *U-*test (**E**). Scale bars: 15 µm (**A**); 50 µm (**C, D**).

### Ambra1 is upregulated in RPE cells after SI treatment and in the macular region of AMD patients

ARPE-19 cells are an RPE-derived human cell line [41] that has been widely used to study the effects of SI *in vitro*. Treatment of ARPE-19 cells with SI for 24 h resulted in loss of cell viability and mitochondria membrane potential, decreased mitochondrial mass, and increased lipid peroxidation (Fig. S6A, B), a similar phenotype to that observed in the RPE of old Ambra*1^+/gt^* mice. Next, we analyzed a dataset derived from transcriptionally-profiled ARPE-19 cells treated for 24 h with SI [42], and found that mRNA expression of *AMBRA1* as well as many genes of the autophagy-initiation complex, including *BECN1*, *ULK1*, and *RB1CC1,* were significantly upregulated in response to SI-induced damage (Fig. S6C, D). Minor increases in mRNA levels were observed for autophagy core genes such as *MAPLC3B*, *ATG5*, *ATG7,* and *ATG16L1* (Fig. S6D). These results suggest upregulation of autophagy machinery to protect against SI-induced damage in RPE cells. Together, these data indicate that *Ambra1-* associated autophagy in RPE may play a key protective role in preventing age- or stress-dependent RPE dysfunction.

Our data show that *Ambra1* haploinsufficiency results in RPE alterations and retinal degeneration that replicates some of the features of macular degeneration, including RPE hypertrophy [43, 44], increased oxidative stress [45], and inflammation [46]. Using a publicly available dataset of RNA sequencing (RNA-seq) profiles of macular and non-macular regions of the human retina and RPE from healthy controls and AMD patients [47], we investigated the expression of the main autophagy regulators (Fig. 9A). While *AMBRA1* mRNA expression was increased in the macular region of AMD donors, no changes were observed in the unaffected non-macular region (Fig. 9B). A similar trend was observed for *RB1CC1* and *ULK1* (Fig. 9C). This gene expression pattern demonstrates that in AMD the autophagy initiation machinery is specifically induced in the macular region, but not in the non-macular region, and that this may constitute an important mechanism to combat the disease.

**Figure 9:**
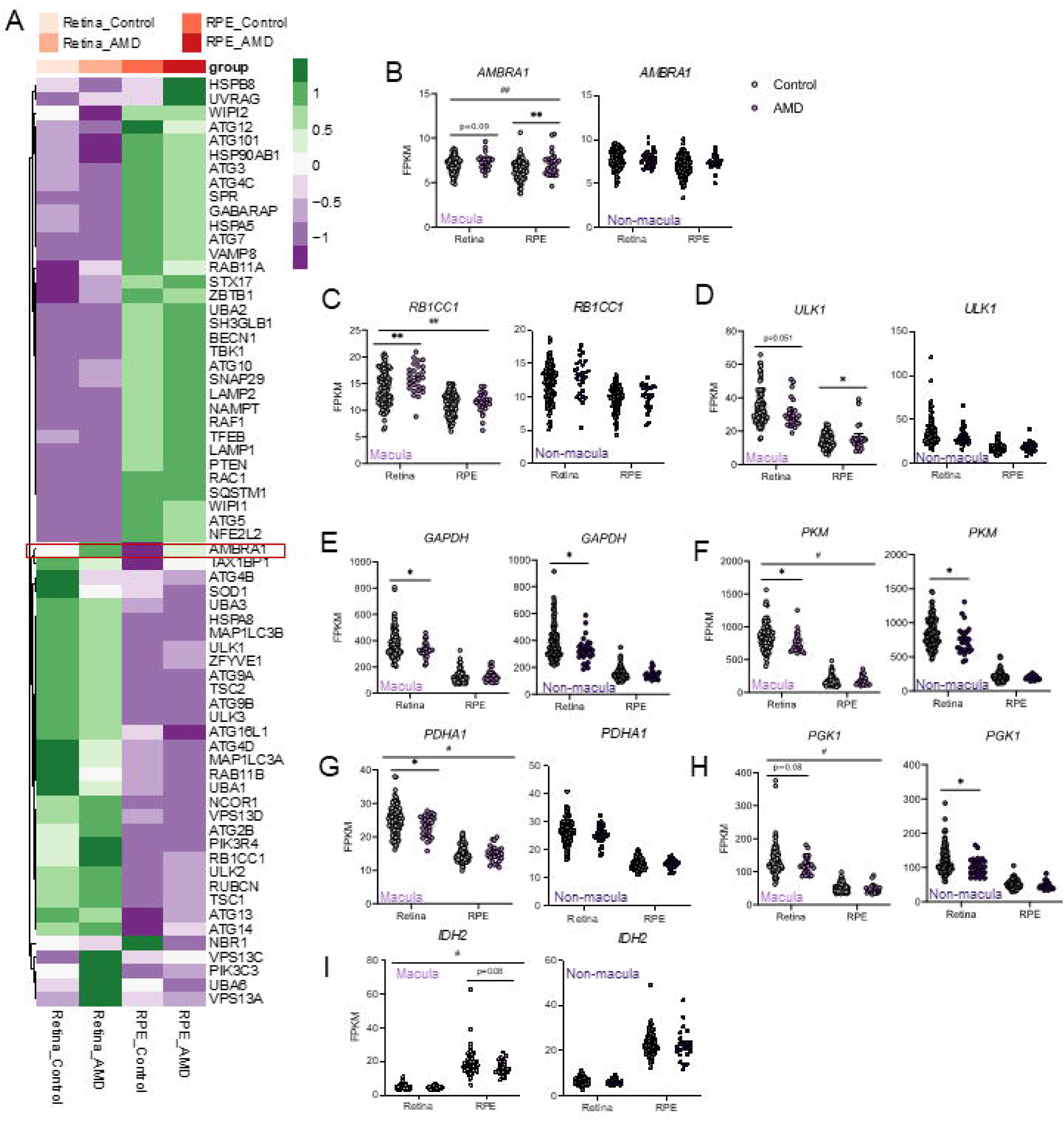
AMD is associated with upregulation of genes of the autophagy initiation machinery and downregulation of glycolysis-related genes. (**A**) Heatmap showing expression analysis of a manually-curated autophagy gene list in the macular region of the retina and RPE of control and AMD patient samples from the GSE135092 dataset (n = 26–105). (**B**) Detailed expression analysis of *AMBRA1* from the aforementioned dataset in macular and non-macular regions (n = 26–105). (**C–J**) Detailed expression analysis of autophagy-related genes: *RB1CC1 (**C**), ULK1 (**D**), GAPDH (**E**), PKM (**F**), PDHA1 (**G**), PGK1 (**H**), IDH2 (**J**)* in the aforementioned dataset in in macular and non-macular regions (n = 26–105). Data are presented as the mean ± SEM. *p<0.05, **p<0.01: two-tailed Student’s *t*-test (B–J). ^#^p <0.05, ^##^p <0.01: two-way ANOVA followed by Fisher’s LSD *post hoc* test for disease (**B–C**, **F–J**).

Metabolic underperformance in the RPE-retina induced by *Ambra1* deficiency was associated with RPE degeneration (Fig. 7 A-B) and loss of visual function (Fig. 2). We therefore investigated whether the aforementioned RNA-seq dataset [47] revealed metabolic alterations in the retina and RPE in AMD patients. In line with our findings in *Ambra1*-deficient retinas, mRNA expression of many glycolytic enzymes was significantly decreased in the retinas of AMD patients; both in macular and non-macular regions, but no changes were observed in the RPE (Fig. 9E-H). These observations, together with the observed decrease in RPE mRNA expression of IDH2 (Fig. 9J), the limiting enzyme in the reductive carboxylation pathway essential for RPE metabolism, suggest decreased mitochondrial oxidative function in the RPE in AMD patients. Taken together, these findings support the view that defective mitochondrial function in the RPE results in a glucose metabolism imbalance, leading to decreased glycolysis in the retina. Moreover, the transcriptional upregulation of autophagy machinery, including *AMBRA1*, observed in the affected macular region in AMD patients suggests a key role of Ambra1-dependent autophagy in protecting the retina in AMD patients.

Taken together, these data demonstrate that *Ambra1* deficiency alters the retina-RPE ecosystem leading to RPE cell death, metabolic underperformance, loss of visual function, and increased susceptibility to AMD-like stress. Our data also show that *Ambra1* expression is selectively upregulated in the macular, but not the non-macular, region of the RPE in AMD patients. Overall, our study demonstrates that AMBRA1*-* mediated autophagy may be a key process for preventing age- and stress-dependent RPE dysfunction and retinal degeneration.

## Discussion

In this study we show that a moderate reduction in autophagy, achieved through monoallelic deletion of *Ambra1* (*Ambra1^+/gt^* mice), exacerbates the retinal degeneration and visual function impairment associated with the aging process. We also describe metabolic alterations associated with mitochondrial dysfunction in *Ambra1^+/gt^* retinas that exacerbate age-dependent photoreceptor degeneration and vision loss. In the RPE, proteostasis alterations were already evident in the young *Ambra1^+/gt^* mice, resulting in RPE atrophy, increased lipid peroxidation, and the formation of lipofuscin aggregates later in life. Upregulation of *Ambra1* and of genes of the autophagy initiation machinery was also observed in the macular, but not the non-macular, region in samples from AMD patients, highlighting the protective role of autophagy in this context. Lastly, *Ambra1* haploinsufficiency sensitized mice to RPE-targeted damage and subsequent photoreceptor death caused by *in vivo* sodium iodate administration, a classical model of RPE degeneration.

*Ambra1^+/gt^* mice provide a useful means of investigating how a slight reduction in autophagy, similar to which accompanies aging, impacts the metabolism and function of the retina and the RPE. As a scaffold protein, AMBRA1 is involved in a wide-range of cellular processes, including the cell cycle [48–51], tumour growth and invasion [52], cell death [53], and gene expression regulation [54]. However, most of these AMBRA1 functions occur in mitotically active tissues or cultured cells. In the retina, which is a post-mitotic tissue, AMBRA1’s contribution to the autophagic pathway may be of greater relevance. In line with this view, we observed no differences in c-MYC, CYCLIN D1, or FOXO3/FOXP3 expression between *Ambra1^+/gt^* mice and control littermates at any age (Fig. S7), suggesting that the observed phenotype may be specifically attributed to the pro-autophagy role of *Ambra1*.

In this study, we use an outbred strain and heterozygous animals, in which some level of autophagic flux is retained. Partial loss of autophagic flux ensures a physiologically relevant context that cannot be achieved using other autophagy-deficient models, which are either perinatal lethal or, in the case of *Atg5* heterozygous animals, retain normal autophagy levels. Also, because AMBRA1 is only implicated in canonical autophagy, in contrast to ATG5 and ATG7 [55], our experimental setting eliminates the confounding effect of LC3-associated phagocytosis in photoreceptor outer segment digestion [16] and of general lysosomal deficiency [56]. Our mouse model therefore provides the unique context in which to examine how the retina-RPE ecosystem copes with the slight reduction in canonical autophagy observed during physiological aging.

The retina’s energy consumption is among the highest of all tissues, even exceeding that of the brain. The retina’s metabolic requirements are fulfilled by glucose, 80% of which is oxidized to lactate via aerobic glycolysis. These reactions produce ATP but also supply many of the building blocks, such as glyceraldehyde 3-phosphate, needed to fuel lipid biosynthesis. This is key to preserve photoreceptor and retinal function [57], as approximately 10% of POS are shed, phagocytized, and degraded daily by the RPE. Carbons from glucose also are completely oxidized to ATP in the mitochondria, as evidenced by the retina’s high level of oxygen consumption [57], and TCA cycle intermediates are also used to support anabolic reactions. These high metabolic requirements of photoreceptors are sustained by the choroidal vasculature, which delivers glucose via the RPE. The RPE is specialized in a form of energy metabolism called reductive carboxylation, which can support redox homeostasis and spares glucose for delivery to photoreceptors [58]. Our data show decreased glycolytic metabolism in the *Ambra1^+/gt^* retina, as demonstrated by reduced mRNA expression of *Hk2*, *Pkm,* and *Gapdh* and an increased pyruvate:lactate ratio. Reduced lactate levels increase the glucose demand of the RPE, as evidenced by the slight increase in GLUT1 expression, thereby decreasing glucose delivery to the retina. Making RPE cells more glycolytic induces degeneration of neighbouring photoreceptors [34, 59]. In normal conditions, the retina would manage this metabolic stress by activating autophagy [60] to cope with starvation and preserve visual function, as observed in mouse models in which autophagy is selectively blocked in the RPE alone [15–17]. However, when autophagy is defective, as in *Ambra1^+/gt^* mice, the retina cannot appropriately respond to starvation by activating autophagy, thus resulting in retinal degeneration and vision defects in the long term. Mitochondrial damage and reduced glucose oxidation also result in a reduction in TCA cycle intermediates and impaired visual function [61], consistent with the marked reductions in amino acids and TCA cycle intermediates that we observed in middle-aged *Ambra1^+/gt^* retinas.

Our findings indicate that marked metabolic alterations in the retinas of middle-aged *Ambra1^+/gt^* mice impair retinal function. Purine metabolism was one of the processes in which the most significant alterations were observed. Although ATP levels were only slightly decreased, we observed a reduction in the total energy charge, in agreement with findings in cancer cells in which autophagy provides metabolic substrates to maintain energy charge and nucleotide pools [35]. Both *de novo* purine biosynthesis and the purine salvage pathway require phosphoribosyl pyrophosphate (PRPP) synthesis from ribose-5-phosphate via the pentose phosphate pathway (PPP) [62], which is highly ATP-consuming. Increases in non-phosphorylated purines and ribose-5-phosphate in old *Ambra1^+/gt^* retinas suggest that altered recycling of nucleotides in autophagy-deficient conditions may enable a compensatory increase in purine biosynthesis, which is reduced due to energy deficit. PPP is also a source of reducing power in the form of NADPH, and we detected an increase in the NADPH:NADP ratio in old *Ambra1^+/gt^* retinas. Another possible explanation is a reduction in the consumption of NADPH for POS lipid synthesis [63, 64]. Our observations suggest a pronounced bioenergetic imbalance in *Ambra1^+/gt^* retinas, in which we also observed an increase in the NADH:NAD ratio. Because NADH is mainly consumed through OXPHOS, we speculated that this finding could reflect dysfunction at the level of the mitochondrial electron transport chain. Notably, mitochondrial mass was unchanged in young *Ambra1^+/gt^* mice, although mitochondrial membrane potential was reduced. In middle-aged *Ambra1^+/gt^* retinas, we observed downregulation of the NAD(P)H quinone dehydrogenase I (*Nqo1*) gene, and of the transcription factors Nrf1 and Nrf2, which drive OXPHOS transcription genes, indicating lower mitochondrial metabolism. Overall, these results suggest alterations in the electron transport chain in the *Ambra1^+/gt^* retina. Recently, it was reported that *Ambra1* mediates PINK1 stabilization during mitophagy [65], leading us to speculate whether mitochondrial alterations observed in *Ambra1^+/gt^* mice could be the consequence of impaired mitophagy.

Our findings in *Ambra1^+/gt^* retinas are also consistent with RPE damage together with alterations in proteostasis that are already evident in young animals. The RPE is a postmitotic tissue that requires efficient quality control mechanisms. Because the RPE is exposed throughout life to oxidative stress as a consequence of the visual cycle [45], autophagy plays an essential housekeeping role and facilitates the removal of intracellular debris derived produced by chromophore recycling [66]. Decreased autophagy has important consequences, resulting in cell enlargement and disruption of the RPE monolayer. These features can be observed in physiological aging in some geriatric mice [38], but are also seen in RPE samples from aged human patients and those with age-related macular degeneration [43]. We propose that *Ambra1* haploinsufficiency gives rise to RPE dysfunction that renders the RPE unable to sustain its metabolic load, ultimately leading to accelerated aging in the neural retina and vision loss. Furthermore, we show that middle-aged *Ambra1^+/gt^* mice are more sensitive to SI, which is commonly used to model RPE degeneration [39].

In middle-aged *Ambra1^+/gt^* mice, cell degeneration occurs earlier in the RPE than the retina, and is characterised by increased inflammation and oxidative stress, features commonly observed in macular degeneration [45]. Indeed, deletion of other autophagy-related genes has been previously reported to induce RPE degeneration [15,56,67,68]. Our analysis of a publicly available RNA-seq dataset of retinal and RPE tissues from control and AMD patients revealed that genes of the autophagy initiation machinery, including *AMBRA1,* were markedly upregulated in the macular region of the RPE compared with unaffected tissue. This suggests that the Ambra1-dependent autophagy initiation complex may be induced to counteract the stress that causes macular degeneration. Glycolysis enzymes were highly expressed in the retina and significantly downregulated in AMD, in line with our findings in *Ambra1*-deficient mice. Taken together, these results reveal the metabolic alterations that occur in the RPE-retina in macular degeneration and highlight the potential role of autophagy induction in preserving retinal homeostasis.

Our findings indicate that maintaining autophagy is crucial to preserve metabolism and retinal function and that slight alterations in autophagy similar to those observed during normal aging promote early RPE damage. Disruption of retinal-RPE homeostasis has important consequences for visual function later in life, as also seen in AMD-like pathologies. Therefore, interventions designed to preserve autophagy could be of therapeutic benefit to prevent blindness in the elderly as well as macular degeneration and other age-dependent retinal diseases.

## Methods

### Animal procedures

All animal experiments were performed following European Union guidelines and the ARVO Statement for the Use of Animals in Ophthalmic and Vision Research. Animal procedures were authorized by the CSIC ethics committee and the Comunidad de Madrid (PROEX244/17). *Ambra1* mutant mice (*Ambra1^+/gt^*) bred on a CD1 background were kindly provided by F. Cecconi [18]. In this study we used male and female *Ambra1^+/gt^* mice and *Ambra1^+/t^* littermates aged 3–4 (young), 12–15 (middle-aged), and 22–26 (old) months. *Ambra1^gt/gt^* mice were not used as these display embryonic lethality [18]. Mice were maintained at the CIB animal facility in a temperature-controlled barrier facility on a 12-h light/dark cycle, with free access to water and food. All animals were sacrificed by cervical dislocation between 9:00 a.m. and 10:00 a.m. to avoid the confounding effects of circadian rhythm changes on autophagy and retinal metabolism.

Tail bud tissue was processed to genotype *Ambra1^+/gt^* animals. Total genomic DNA was isolated using the NZY Tissue gDNA Isolation kit (#MB13503; Nzytech). Polymerase chain reaction (PCR) was performed using the primers 5’-CCCAGTCACGACGTTGTAAAA-3’ (primer A) and 5’-TCCCGAAAACCAAAGAAGA-3’ (primer B), mapping downstream and upstream of the gene-trap insertion site corresponding to the lacZ reporter sequence. PCR conditions were as follows, for a total of 35 cycles: unfolding temperature, 95°C for 2 min; annealing temperature, 58°C for 54 s; elongation temperature, 72°C for 45 s.

For the SI model, 6.25 mg/mL of sterile sodium iodate (SI, [NaIO3]) solution was freshly prepared from solid SI (S4007; Sigma-Aldrich) diluted in PBS (D8537; Sigma-Aldrich). Mice received a single intraperitoneal injection of 50 mg/kg of SI solution (experimental group) or vehicle (control group) and were sacrificed 1 week after injection.

### Electroretinogram recordings

For ERG recordings mice were first allowed to adapt to darkness overnight (o/n), and then subsequent manipulations were performed in dim red light. Mice were anesthetized by i.p. injection with ketamine (95 mg/kg) and xylazine (5 mg/kg) solution while placed on a heating pad set at 37°C. Pupils were dilated with a drop of 1% tropicamide (Colircusi Tropicamida; Alcon Cusi). A drop of 2% Methocel (Hetlingen) was placed in each eye immediately before placement of the corneal electrode to optimize electrical recording.

Flash-induced ERG responses to light stimuli produced with Ganzfeld stimulator were recorded in the right eye. A photometer was used to measure light intensity at the level of the eye (Mavo Monitor USB). In scotopic conditions, 4–64 consecutive stimuli were averaged with an interval of 10 s for dim flashes and 60 s for the highest intensity flashes. In photopic conditions, the interval between light flashes was set at 1 s. ERG signals were band-filtered between 0.3 and 1000 Hz and amplified with an amplifier (CP511 AC amplifier; Grass Instruments). A power laboratory data acquisition board (AD Instruments) was used to digitize electrical signals at 20 kHz. An electrode (Burian-Allen electrode; Hansen Ophthalmic Development Laboratory) was fixed on the corneal lens and a reference electrode was placed in the mouth, with a ground electrode attached to the tail to perform bipolar recordings. In dark-adapted conditions, the following responses were recorded: rod responses to light flashes of -2 log cd·s·m^-2^; and mixed responses to light flashes of 1.5 log cd·s·m^-2^. White flashes of -1.5 log cd·s·m^-2^ were used in a recording frequency range of 100–10,000 Hz to isolate the oscillatory potential (OP). In light-adapted conditions, cone-mediated responses to light flashes of 1.5 log cd·s·m^-2^ were recorded on a rod-saturating background of 30 cd·m^-2^. Wave amplitudes of scotopic rod responses (b-rod), OP, mixed responses (a-mixed and b-mixed), and photopic cone responses (b-phot and flicker) were measured off-line by an observer blind to the experimental condition of the animal. Also, the b:a wave ratio was calculated to better understand the electroretinographic responses.

### Tissue preparation for imaging

For retina and RPE/choroid flat mounts, eyes were enucleated and briefly washed in ice-cold PBS. The optic nerve, cornea, and lens were gently removed and the resulting posterior eyecup was fixed in freshly prepared ice-cold 4% (w/v) PFA (171010, EMS) in PBS for 30 min. Four perpendicular incisions were made and the retina and RPE/choroid were gently separated and fixed for an extra 1.5 h at room temperature (RT). Tissues were kept in 0.01% azide in PBS at 4°C.

For cryosections, mouse eyeballs were fixed o/n in 4% (w/v) PFA (171010; EMS) at 4°C. Next, eyeballs were cryoprotected and embedded in OCT (Tissue Tek, Sakura Finetek). Sections (12 μm) were cut on a cryostat (Leica Microsystems).

### Ex-vivo analysis of autophagic flux

Autophagic flux was assessed as previously described [69]. Dissected retinas werewas placed (photoreceptors down) in Millicell support inserts (Millipore) and maintained in DMEM (Dulbeccós Modified Eagle Medium; 41966-029, Gibco) with 1% glutamine (2 mM; 25.030 Gibco), 1% penicillin-streptomycin (0.5 mg/mL; 11568876, Gibco), and 1 μM insulin (I2643, Sigma) at 37°C in a 5% CO^2^ atmosphere. Retinas were treated for 3 h with 100 μM leupeptin (L-2884-50, Sigma) and 20 mM ammonium chloride (A9434-500, Sigma) to inhibit lysosomal proteolysis. No cell death was observed during culture and there were no differences in the LC3-II:LC3-I ratio between freshly isolated retinas and *ex vivo* retinal cultures (data not shown). After culture, retinas were stored dried at -80°C.

### Cell culture

ARPE-19 cells (ATCC, CRL-2302) were grown in DMEM: F12 (1:1) supplemented with 15% FBS, 1% glutamine (2 mM), and 1% penicillin-streptomycin (0.5 mg/mL). Cells were seeded at a density of 10^5^ cells/mL in 24-well plates. For immunofluorescence, cells were grown on 12-mm glass coverslips and immunostained as previously described. For flow cytometry, cells were incubated for 30 minutes with DiOC6(3) and Mitotracker Deep Red (MTDR), DAPI was added to select the viable population by exclusion gating, and at least 10000 events were recorded using a CytoFlex S system (Beckman Coulter).

### Immunofluorescence

Retinal or RPE/choroid samples were permeabilized with 2% or 0.2% (v/v) Triton X-100 and blocked for 1 h with BGT (3 mg/mL BSA, 0.25% Triton X-100, 100 mM glycine in PBS) or block/perm solution (10% normal goat serum, 0.1% Triton X-100 in PBS), respectively. Retinal cryosections were re-fixed with 4% (w/v) PFA for 15 min, washed in PBS, and permeabilized with Triton X-100 1% (v/v) for 1 h. Next, sections were blocked for 1 h with BGT. Samples were incubated with primary antibodies o/n at 4°C. The following primary antibodies were used: anti-LC3 (Novus, NB100-22020, 1/200), anti-ConeArrestin (Millipore, AB15282, 1/200), anti-PKCα (Sigma, P4334, 1/200), anti-RG opsin (Millipore, AB5405, 1/200), anti-BRN3A (Millipore, MAB1585, 1/100), anti-GFAP (DAKO, Z0334, 1/500), anti-GS (Millipore, MAB302, 1/500), anti-IBA1 (Wako, 019-19741, 1/500), anti-4-HNE (Abcam, ab46545, 1/100), anti-TOMM20 (Santa Cruz Biotechnology, sc-11415, 1/200), anti-LAMP2 (ABL93, DSHB, 1/200), anti-GAPDH (Abcam, ab8245, 1/100), anti-HK2 (C64G5, ab 2867, Cell Signaling), and anti-GLUT1 (LAB VISION, Fremont, CA, 1/100). After washing with PBS, tissues were incubated for 1 h at RT in darkness with the secondary antibodies (1/200; Alexa 488, Alexa 568 and Alexa 647; Invitrogen) and DAPI (4’,6-diamino-2-phenylindole) (D9542; Sigma; 1 μg/mL). Additionally, RPE/choroid flat mounts were counterstained with phalloidin (A12380; Invitrogen, 1/500) together with the secondary antibodies. Finally, flat-mounted retinas were mounted with Fluoromount (100-01; Bionova) between 2 sealed coverslips. Cryosections were mounted with DABCO (1,4-diazabicyclo[2.2.2]octane) (D27802; Sigma) and sealed with nail polish. The specificity of the LC3 antibody was verified by complete colocalization of anti-LC3 immunostaining and GFP-LC3 reporter in ARPE-19 cells (data not shown).

The ProteoStat® assay (ENZ-51023-KP050; Enzo) was performed following the manufacturer’s instructions. Cryosections were incubated with the dye at 1/500 dilution for 1 h at RT in a wet chamber before DAPI counterstaining. TUNEL assay (DeadEnd™ Fluorometric TUNEL System; #G3250; Promega) was used to detect apoptotic cells in cryosections following the manufacturer’s directions. Briefly, once the primary antibody was washed, cryosections were incubated for 30 min with TUNEL buffer, after which the TUNEL reaction (1.9% TdT, 9.8% dNTPs, and 88.3% TUNEL buffer) was performed in darkness for 1 h at 37°C. Finally, saline-sodium citrate (SSC) provided in the kit (20X) was diluted to a concentration of 2X and added to cryosections to stop the TUNEL reaction.

Confocal z-stacks were obtained with Leica TCS-SP5-A0BS or Leica TCS SP8 STED 3X microscopes (Leica Microsystems). To assess retinal layer thickness, images of DAPI-stained cryosections were acquired at 40X with a Leica DMI6000B fluorescence microscope coupled to a Leica AF6000 LX multidimensional system (Leica Microsystems).

Autofluorescence was assessed using single-channel lambda-scan (xyλ) with a Leica TCS SP8 STED 3X microscope (Leica Microsystems). Briefly, cryosections were mounted using DABCO mounting medium, stimulated using the UV-laser line (λex=405 nm) and autofluorescence was captured in 10-nm steps (λem=420–740 nm). The RPE was manually delimited and mean fluorescence intensity was measured.

### Image analysis and data quantification

Unless stated otherwise, maximum projections of all z-stacks are displayed in representative images (z-step: 0.5 and 1 µm for RPE and retina, respectively; 1 μm for cryosections). For retinal thickness measurements, each retinal layer was manually measured using the straight-line tool in ImageJ. Quantification of mean fluorescence intensity (MFI) was performed using unprocessed images at maximal projection. Positive cells for a given marker were determined by manual counting plane by plane in a given z-stack. LC3 puncta were quantified using a manually designed Fiji-based plugin, which takes into account the 3D component. Briefly, a fluorescence intensity threshold is set to discriminate positive signal from background. Next, using a minimum voxel size threshold, the 3D objects counter tool is used to detect the number of objects in a 3D confocal z-stack. Finally, the number of puncta obtained is corrected to the thickness of the retina based on the z-stack size.

All confocal images from the same experiment were acquired using the same laser intensity and photomultiplier settings to avoid any variability or bias. Moreover, only nuclear DAPI staining was used to select the field to photograph. At least 3 sections per mice and 4 retinal regions were included in each analysis.

### Quantitative RT-PCR

Retinal RNA was extracted using TRIzol reagent (Invitrogen) and converted into cDNA using the High-Capacity cDNA Reverse Transcription Kit (4374966; Applied Biosystems) following the manufacturer’s instructions. Quantitative reverse transcription PCR (RT-PCR) was performed with 10 ng of cDNA in a Light Cycler^®^ Instrument (Roche) using LightCycler^®^ 480 probes master mix (4887301001, Roche) and Taqman probes (Life Technologies). The following TaqMan gene expression probes were used: *Sqstm1* (Mm00448091_m1), *Atg5* (Mm00504340_m1), *Wipi2* (Mm00617842_m1), *Tfeb* (Mm00448968_m1), *Ppargc1a* (Mm01208835_m1), *Nrf1* (Mm01135606_m1), *Nfe2l2 (Mm00477784_m1), Tfam* (Mm00447485_m1), *Timm23* (Hs00197056_m1), *COX IV* (Mm01250084_m1), *Gapdh* (Mm99999915_g1), *Hk2* (Mm00443385_m1), *Pkm* (Mm00834102_gH), *Nqo1* (Mm01253561_m1) and *Pdha1* (Mm00468675_m1). Duplicates were included in the analysis and results were normalized to expression levels of 18s RNA (Hs99999901_s).

### Western blot

Retinal protein extracts were obtained in lysis buffer consisting of 50 mM Tris-HCl (pH 6.8), 2% SDS (w/v), 10% glycerol (v/v), phosphatase inhibitors (1 mM sodium fluoride [201154; Sigma-Aldrich], 1 mM sodium orthovanadate [S6508; Sigma-Aldrich], 5 mM sodium pyrophosphate [21368; Sigma-Aldrich], and protease inhibitor cocktail [P8783; Sigma-Aldrich]). Isolated retinas were homogenized using a plastic pestle until they were completely disaggregated. Next, samples were heated for 12 min at 95°C and stored at -20°C. Pierce BCA Protein Assay Kit (23227, Pierce Thermo Fisher Scientific) was used to determine protein concentration following the manufactureŕs instructions.

Total protein extract (15 μg) was mixed with 10 mM DTT and bromophenol blue and loaded into Any kD Criterion TGX Precast Gels (567-1124, Bio-Rad). PVDF membranes (170-4157, Bio-Rad) were then activated with 100% methanol and total protein extracts were transferred for 14 min at 25 V using a Trans-Blot Turbo Transfer System (Bio-Rad). Membrane-bound proteins were visualized with Ponceau S (78376; Sigma) in 5% acetic acid (1.00063.1000; Merck). Membranes were blocked for 1 h at RT with 5% non-fat milk in PBS-T (1X PBS, 0.5% Tween 20 [v/v]) and incubated o/n at 4°C with the following primary antibodies at a dilution of 1/1000: anti-LC3 (Sigma, L7543), anti-Atg5-12 (Abnova, PAB12482), anti-FoxO3a (Abcam, ab70315), anti-Foxp3(Biolegend, 320007), anti-c-myc (Abcam, ab32072), anti-cyclinD1 (Santa Cruz Biotechnology, sc-718), anti-Tim23 (BD Biosciences, 611222), anti-Tomm20 (Santa Cruz Biotechnology, sc-11415), γH2AX (Millipore, 07-164), anti-ubiquitin (P4D1, Santa Cruz Biotechnology, sc-8017), anti-tubulin (T6199; Sigma) anti-vinculin (Abcam, ab129002) and anti-Gapdh (Abcam, ab8245). After washing with PBS-T, membranes were incubated with the secondary antibody coupled with horseradish peroxidase (1/2000). Finally, membranes were revealed using the HRP Pierce ECL Western Blotting substrate (32106; Thermo scientific) and film (AGFA) after exposure for varying durations. Band densitometry analysis was performed using Quantity One 1-D Analysis Software (Bio-Rad). To normalize across different gels and to encompass all 3 ages and genotypes, a sample from the ARPE-19 cell line was included in all 3 gels as an internal loading control.

### Flow cytometry of dissociated retinas

Retinas were dissected and incubated with 1% (w/v) trypsin (25300054; Gibco) in DMEM for 5 min at 37°C. Each sample was centrifuged at 200 × g for 5 min and pellets were incubated with 40 nM DiOC6 (D273, Invitrogen) for 15 min at 37°C and 5% CO_2_ to assess mitochondrial membrane potential. Finally, dissociated retinas were resuspended in 300 μL of DMEM and analyzed in an XL flow cytometer (Beckman Coulter).

### Mass spectrometry and bioinformatics analysis

Metabolomic analyses were conducted using mass spectrometry as previously described [33]. Briefly, retinal tissue (20 mg) was obtained from 2 pools of 4 retinas from 4- and 12-month-old *Ambra1^+/gt^* mice and *Ambra1^+/+^* littermates. For each condition, tissue was weighed and solubilized with 500 μL of cold (−20°C) lysate buffer (MeOH/water/chloroform [9/1/1] with internal standards) in 1.5 mL polypropylene (Precellys lysis tubes). Tissue was homogenized 3 times for 20 s using a Precellys 24 tissue homogenizer (Bertin Technologies). The homogenized tissue was centrifuged (10 min at 15,000 × g and 4°C) and the upper phase of the supernatant was collected. A second round of extraction was performed using 500 μL of lysate buffer, followed by homogenization and centrifugation to collect the supernatant, which was added to the first supernatant. All supernatant collected was then evaporated in microcentrifuge tubes at 40°C in a pneumatically assisted concentrator (Techne DB3). Methanol (100%, 300 μL) was added to the dried extract and the resulting solution split into 2 volumes of 150 μL each. The first part was used for the GC-MS (gas chromatography-mass spectrometry), and the second for LC-MS (liquid chromatography-mass spectrometry) analysis. The GC-MS aliquots were solubilized in methanol and transferred to a glass tube. Next, 50 μL of methoxyamine (20 mg/mL in pyridine) was added to the dried extracts, which were stored in darkness for 16 h at RT. The following day, 80 μL of N-methyl-N-(trimethylsilyl)trifluoroacetamide (MSTFA) was added to perform the final derivatization at 40°C for 30 min. Samples were then transferred to glass vials and directly injected into the GC-MS system. After the second evaporation of the LC-MS aliquots, the LC-MS-dried extracts were solubilized in 50 μL of Milli-Q water, centrifuged (10 min at 15,000 × g at 4°C), transferred to glass vials, and injected into the UHPLC/MS (ultra-high-performance liquid chromatography/mass spectrometry) system. Target analysis was performed as previously described [70]. Manual verifications and QC protocols were followed to select 83 metabolites for GC targeted analysis and 17 metabolites for LC targeted analysis.

After the metabolomic analyses, a peak area proportional to concentration was obtained for each metabolite. Based on these data, metabolite ratios were calculated as the coefficient between arbitrarily chosen metabolites. Principal Component Analysis (PCA) was performed using prcomp function in R. Raw data were processed and analyzed using MultiExperiment Viewer (MEV) software. Skillings-Mack test was performed with significance set at 0.05 to compare metabolite data between *Ambra1^+/gt^* and *Ambra1*^+/+^ littermate replicates. Next, unsupervised hierarchical clustering (HCL) was performed to cluster significantly different metabolites and construct heatmap representations. Finally, the online MetaboAnalyst 4.0 tool (http://www.metaboanalyst.ca/) for metabolomic data analysis and interpretation was used to perform enrichment analysis of differentially obtained metabolites between *Ambra1^+/gt^* and *Ambra1*^+/+^ duplicates.

### Bioinformatics analysis

Processed datasets (GSE142591 and GSE135092) were downloaded from the GEO database. FPKM (fragments per kilobase of transcript per million mapped reads) values were obtained from the already deposited data. From all detected genes in the available datasets, manually-curated autophagy or glycolysis gene lists were subtracted using R programming. Next, using pheatmap in the R package, heatmaps were generated and hierarchical clustering performed for both samples and genes. Finally, statistical analysis and plotting of selected genes was performed using GraphPad Prism software (GraphPad Software, Inc.).

### Statistical analysis

All numeric results are presented as mean values ± SEM. Statistical analyses were performed using GraphPad Prism software (GraphPad Software, Inc.). To test for differences between genotypes, two-tailed Student’s t-tests were used. To test for differences between genotypes and age class two-way analyses of variance (ANOVA) were applied and in the case of a significant interaction, differences between genotypes within each age class were assessed using Fisher’s least significant difference (LSD) post hoc tests. If assumptions of normality and homoscedasticity were not met, we applied non-parametric tests such as Mann-Whitney *U*-test for two-group comparisons and Kruskal-Wallis when more than two groups were compared. Electroretinographic responses were analyzed with general linear models using the lmer function implemented in R. The statistical model contained the electroretinographic response as response variable, genotype and sex as factors, the exact age in number of days as a covariate, the interaction between genotype and number of days, and the experimental group as random factor. Model assumptions were tested. If the normality assumption was violated, transformations were applied and in the presence of heteroscedasticity, weighted least square regressions were performed. For all tests the significance level was set at p<0.05 (two-tailed). The number of animals used per experiment is stated in the corresponding figure legend.

## Abbreviations

AMD: age-related macular degeneration
CNS: central nervous system
FACS: fluorescence-activated cell sorting
GCL: ganglion cell layer
GFAP: glial fibrillary acidic protein
GS: glutamine synthetase
HCL: hierarchical clustering
INL: inner nuclear layer
IPL: inner plexiform layer
LC/GC-MS: liquid chromatography/gas chromatography–mass spectrometry
MA: middle-aged
MDTR: MitoTracker Deep Red
MFI: mean fluorescence intensity
NL: NH_4_Cl and leupeptin
Nqo: NAD(P)H quinone dehydrogenase
ONL: outer nuclear layer
OPL: outer plexiform layer
OP: oscillatory potentials
OXPHOS: oxidative phosphorylation
PCA: principal component analysis
PCR: polymerase chain reaction
PKC: protein kinase C
PPP: pentose phosphate pathway
POS: photoreceptor outer segment
PRPP: phosphoribosyl pyrophosphate
RGC: retinal ganglion cells
RPE: retinal pigmented epithelium
SI: sodium iodate
TCA: tricarboxylic acid; 4-HNE: 4-hydroxynonenal

## Acknowledgments

We thank Francesco Cecconi for the *Ambra1* mice; Daniel Lietha, Aurora Gómez-Durán and Pura Muñoz-Cánoves for reagents; Guido Kroemer for the access to the metabolomics facility at the IGR, Villejuif, France; Owen Howard for English-language editing; and Helena Vieira and Marta Agudo Barriuso for comments on the manuscript. We are also grateful to M. Teresa Seisdedos and Gema Elvira at the Confocal Microscopy facility at the CIB Margarita Salas, Ana Oña Blanco and the technical staff of the Confocal Microscopy facility at the CNB, and the technical staff at the animal facility of the CIB Margarita Salas for their support. Thanks also to Servier Medical Art (https://smart.servier.com/) for the images used in the graphical abstract.

## Figure legends

Supplementary Figure 1. Autophagy is decreased with aging and in *Ambra1^+/gt^* mice.

(**A**) Decreased *Ambra1* mRNA expression as determined by qPCR of whole retina extracts from young, middle-aged, and old *Ambra1*^+/+^ and *Ambra1*^+/gt^ mice (n = 3–4 per group). (**B-C**) Autophagic flux comparing *Ambra1^+/+^* and *Ambra1^+/gt^ ex vivo* retinal cultures treated with protease inhibitors (NL) with untreated retinas (Ctrl) from middle-aged (**B**) and old (**C**) mice. (**D**) Western blot showing age-related changes in the expression of the autophagy proteins Atg5-Atg12 conjugate and unconjugated Atg5 in *Ambra1^+/+^* and *Ambra1^+/gt^* mice, and corresponding quantification (right). (**E**) LC3 staining in different retinal layers. Corresponding quantification of LC3 puncta is shown on the right. OS, outer segments; IS, inner segments; ONL, outer nuclear layer; INL, inner nuclear layer; GCL, ganglion cell layer. Data are presented as the mean ± SEM. *p <0.05, ***p <0.001: two-tailed Student’s *t*-test (**E**); two-way ANOVA followed by post hoc Fischer’s LSD test for genotype (**A**) and age **(E**). Scale bars: 25 µm.

Supplementary Figure 2. The age-associated decline in electrophysiological responses is exacerbated in *Ambra1^+/gt^* mice. (**A–E)** Electroretinographic responses measured in *Ambra1^+/+^* (n = 58) and *Ambra1^+/gt^* (n = 32) mice at different ages. The predicted regression lines and the associated 95% confidence intervals are shown. Significant slope differences are indicated: \**p* <0.05, **p <0.001, \*\*\**p* <0.0001.

Supplementary Figure S3. Morphological alterations and increased inflammation in old *Ambra1^+/gt^* retinas. (**A**) Detection of apoptotic cells (TUNEL, green) and corresponding quantification (right) in young, middle-aged, and old *Ambra1*^+/+^ and *Ambra1*^+/gt^ mice (n = 3–9). (**B**) Immunostaining of red/green cones (RG opsin, cyan; left) and corresponding quantification (right) in temporal to nasal regions in old *Ambra1*^+/+^ and *Ambra1*^+/gt^ mice (n = 5). **(C)** Immunostaining of microglial cells (IBA1, magenta) in retinal flat mounts (left) from old *Ambra1*^+/+^ and *Ambra1^+/gt^* littermates. Two distinct microglial cell morphologies (ramified and ameboid) were observed (right) (n = 5–6 per group). Arrows indicate ameboid microglial cells. (**D**) Representative images showing immunostaining of gliosis (GFAP, magenta) and Müller cells (GS, green) in young, middle-aged, and old *Ambra1^+/+^* (left panels) and *Ambra1^+/gt^* mice (right panels). Nuclei are counterstained with DAPI (grey). Data are presented as the mean ± SEM. \**p* <0.05: two-way ANOVA followed by Fisher’s LSD *post hoc* for genotype (**B**). Scale bars: 50 µm (**A, C, D**); 25 µm (**B**).

Supplementary Figure S4. Detailed metabolomic analysis of *Ambra1^+/gt^* retinas. (**A**) Heat map and hierarchical clustering (HCL) analysis of all metabolites from young and middle-aged (MA) *Ambra1*^+/+^ and *Ambra1^+/gt^* retinas (n = 2 per group). (**B**) Pairwise Pearsońs correlation of biological replicates from a metabolomic study of whole retina extracts from young and middle-aged (MA) *Ambra1*^+/+^ and *Ambra1^+/gt^* mice (n *=* 2 per group). (**C**) Pathway enrichment analysis of metabolites for which significant differences were observed between *Ambra1*^+/+^ and *Ambra1^+/gt^* duplicates (n = 2 per group). Statistical significance (Skillings-Mack test) was set at p <0.05.

Supplementary Figure S5: Middle-aged *Ambra1^+/gt^* mice show alterations in autophagy and metabolism in the RPE. (**A, B**) RPE flat mounts from *Ambra1^+/+^* and *Ambra1^+/gt^* littermates immunostained for LC3 and corresponding quantification (n = 3 per group). (**B**) Quantification of LC3+ puncta in **A**. (**C, D**) Immunostaining for LC3 in RPE cryosections from *Ambra1^+/+^* and *Ambra1^+/gt^* littermates and corresponding quantification of LC3+ puncta in young and middle-aged animals (n = 5 per group). (**E**) Levels of TOMM20 and TIM23 proteins in the RPE were evaluated by Western blot in middle-aged *Ambra1^+/+^* and *Ambra1^+/gt^* mice. (**F**) Immunostaining of GLUT1 in RPE cryosections in young, middle-aged, and old *Ambra1^+/+^* and *Ambra1^+/gt^* littermates. Corresponding quantification of GLUT1 mean fluorescence intensity (n = 3–5 per group). (**G**) Immunostaining of GAPDH in RPE cryosections from young and middle-aged *Ambra1^+/+^* and *Ambra1^+/gt^* littermates. Corresponding quantification of GAPDH in young, middle-aged, and old *Ambra1^+/+^* and *Ambra1^+/gt^* littermates (n = 3–5 per group). (**H**) Immunostaining of microglial cells (IBA1, red) (left) in the RPE and choroid of young and middle-aged *Ambra1^+/+^* and *Ambra1^+/gt^* mice (n = 3–5 per group). Corresponding quantification is shown on the right. Nuclei were counterstained with DAPI (blue or grey). Data are presented as the mean ± SEM. *p <0.05; two-way ANOVA followed by Fisher’s LSD *post hoc* test for genotype (**B, D**); two-tailed Student’s *t*-test (E, TIM23); or Mann Whitney *U*-test (E, TOMM20). Scale bars: 50 µm € and 25 µm (**A**, **F**, **G** and **H**).

Supplementary Figure S6: Sodium iodate (SI) induces cell death and transcription of autophagy machinery genes in ARPE19 cells. (**A**) ARPE-19 cells were treated with 20 mM SI for 24 h and then analyzed by flow cytometry to assess viability (DAPI exclusion assay), ΔΨm (DiOC6(3)), and mitochondrial mass (MitoTracker Deep Red, MTDR). n = 3–4 independent experiments with 2 biological replicates. (**B**) ARPE-19 cells were treated with 20 mM SI for 24 h and lipid peroxidation was assessed by immunofluorescence for 4-HNE (green). Nuclei were counterstained with DAPI (blue). n = 6 biological replicates from 2 independent experiments. (**C, D**) RNA-seq dataset (GSE142591) from ARPE19 cells treated with 20 mM SI for 24 h. (**C**) A manually-curated autophagy gene list was used to generate a heatmap of unsupervised hierarchical clustered samples and genes (n = 3). (**D**) Gene expression based on normalized counts of selected autophagy genes (*AMBRA1, BECN1, ULK1, RB1CC1, MAP1LC3B, ATG5, ATG7, ATG16L1*) from the same dataset (n = 3). Data are presented as the mean ± SEM. *p<0.05, **p<0.01: two-tailed Student’s *t*-test (**A**, **B**, **D**). Scale bar: 50 µM.

Supplementary Figure S7: Non-autophagic functions of Ambra1 are unchanged in the RPE of *Ambra1^+/gt^* mice. (**A**) Protein levels of FOXP3, FOXO3A, CYCLIN D1 and c-MYC were evaluated by Western blot in the RPE in young, middle-aged, and old *Ambra1^+/+^* and *Ambra1^+/gt^* mice. (**B**) Quantification of Western blots shown in A, expressed as protein level relative to that of the loading control (vinculin) (n = 4–5 per age/genotype). Statistical analysis was performed by 2-way ANOVA followed by Fisher’s LSD *post hoc* test. *p<0.05.

Supplementary Figure S8: Graphical abstract. *Ambra1^+/gt^* mice display diminished autophagy activity in both the retina and RPE. This decrease in autophagy is accelerated relative to that which occurs during physiological aging. This age-associated autophagy deficiency results in premature degeneration of the RPE and retina and loss of visual function. Cellular and functional degeneration the RPE-retina is associated with protein accumulation, oxidative stress, and inflammation. Cellular damage disrupts the metabolic balance of the RPE-retina, leading to reduced mitochondrial mass in degenerating RPE cells and mitochondrial dysfunction and reduced glycolysis in the retina. Graphical abstract images templates were obtained from Servier Medical Art (https://smart.servier.com/)

