## Supplementary figures for "Ambra1 haploinsufficiency results in metabolic alterations and exacerbates age-associated retinal degeneration"

### Supplementary Figure S1

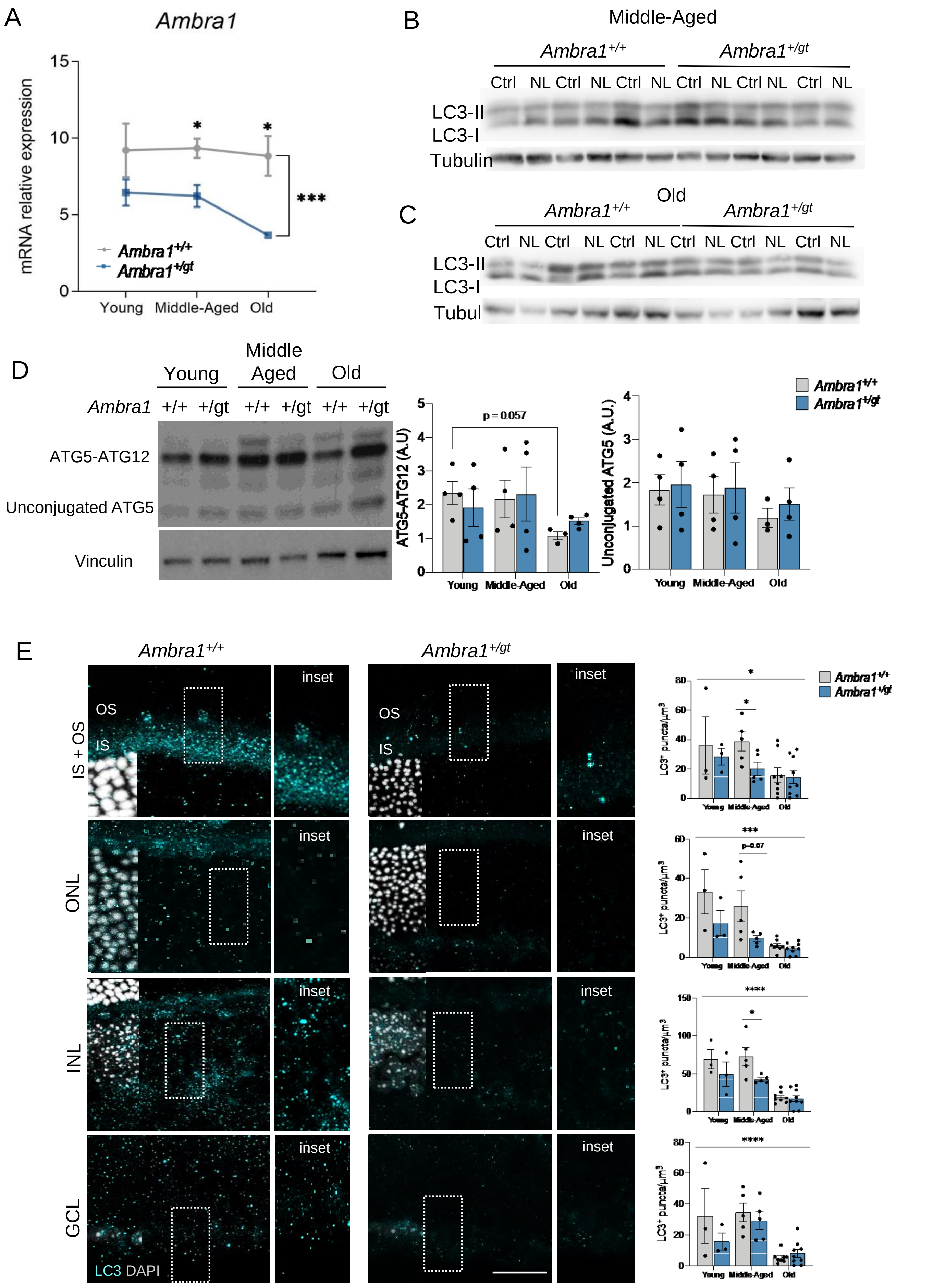

Supplementary Figure 1. Autophagy is decreased with aging and in *Ambra1*$^{+/gt}$ mice. (**A**) Decreased *Ambra1* mRNA expression as determined by qPCR of whole retina extracts from young, middle-aged, and old *Ambra1*$^{+/+}$ and *Ambra1*$^{+/gt}$ mice (n = 3–4 per group). (**B-C**) Autophagic flux comparing *Ambra1*$^{+/+}$ and *Ambra1*$^{+/gt}$ *ex vivo* retinal cultures treated with protease inhibitors (NL) with untreated retinas (Ctrl) from middle-aged (**B**) and old (**C**) mice. (**D**) Western blot showing age-related changes in the expression of the autophagy proteins Atg5-Atg12 conjugate and unconjugated Atg5 in *Ambra1*$^{+/+}$ and *Ambra1*$^{+/gt}$ mice, and corresponding quantification (right). (**E**) LC3 staining in different retinal layers. Corresponding quantification of LC3 puncta is shown on the right. OS, outer segments; IS, inner segments; ONL, outer nuclear layer; INL, inner nuclear layer; GCL, ganglion cell layer. Data are presented as the mean ± SEM. *p <0.05, ***p <0.001: two-tailed Student´s *t*-test (**E**); two-way ANOVA followed by post hoc Fischer's LSD test for genotype (**A**) and age **(E).** Scale bars: 25 µm.

### Supplementary Figure S2

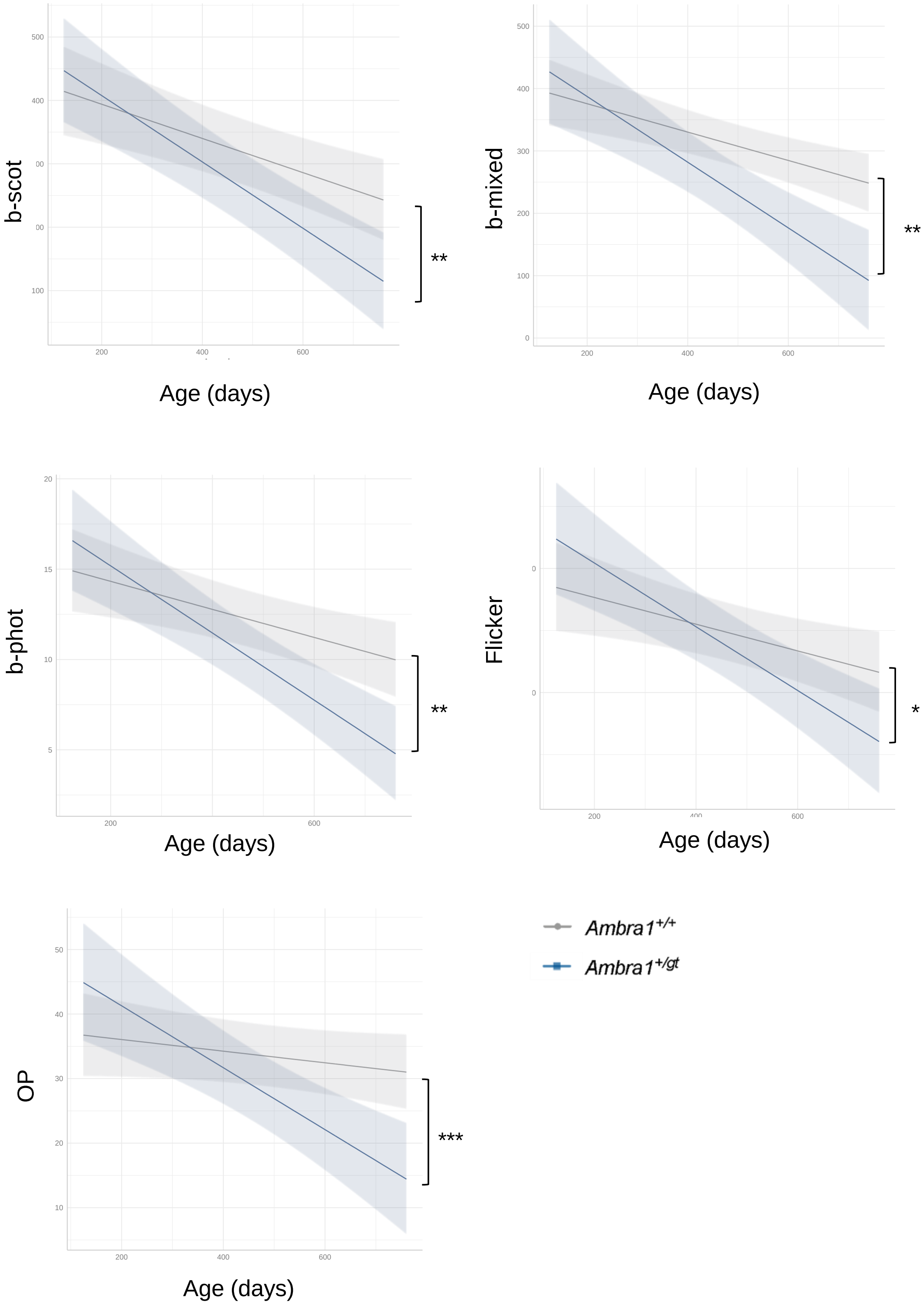


Supplementary Figure 2. The age-associated decline in electrophysiological responses is exacerbated in *Ambra1*$^{+/gt}$ mice. (**A–E)** Electroretinographic responses measured in *Ambra1*$^{+/+}$ (n = 58) and *Ambra1*$^{+/gt}$ (n = 32) mice at different ages. The predicted regression lines and the associated 95% confidence intervals are shown. Significant slope differences are indicated: **p* <0.05, **p <0.001, ****p* <0.0001.

### Supplementary Figure S3

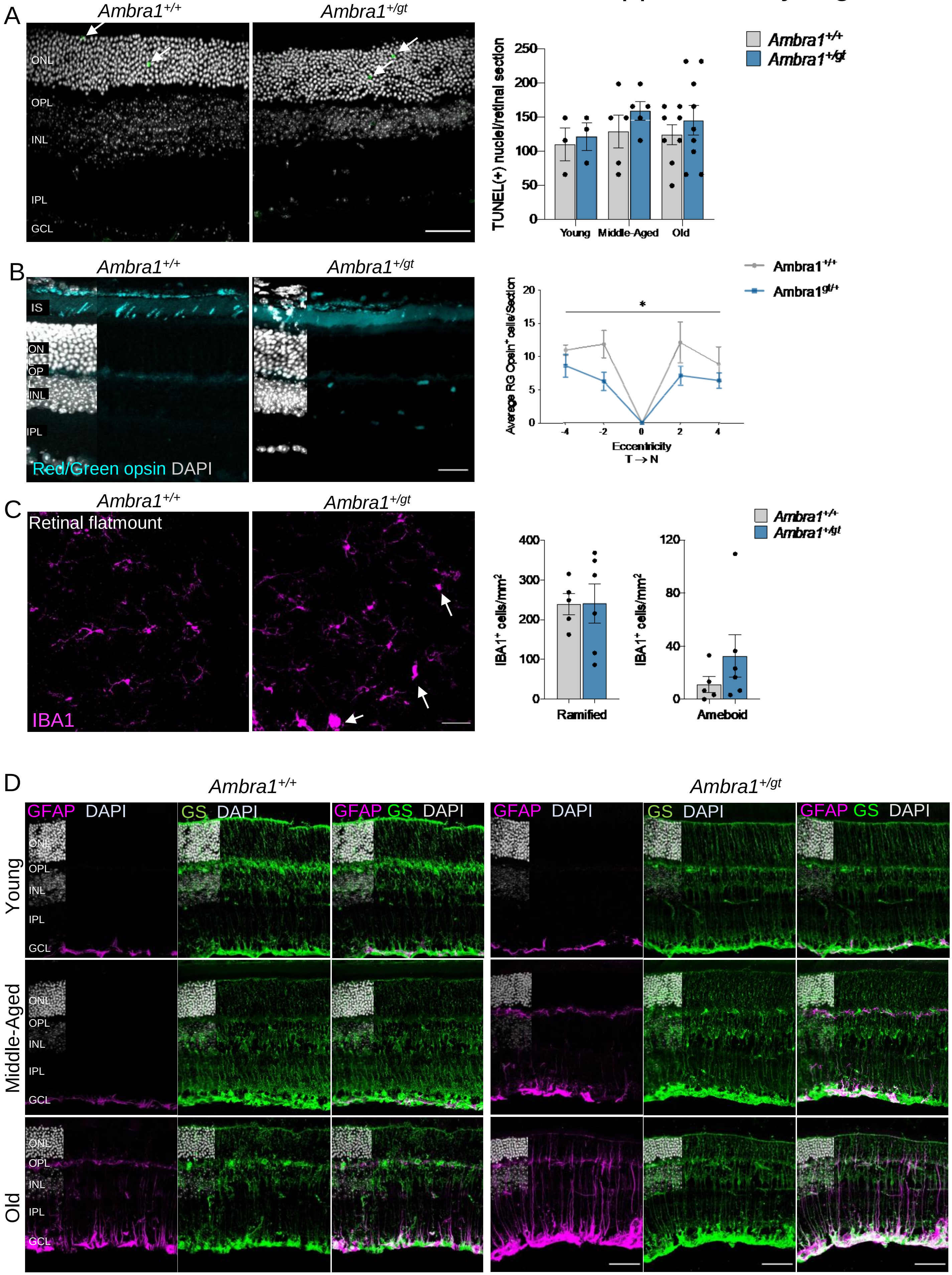

Supplementary Figure S3. Morphological alterations and increased inflammation in old *Ambra1*$^{+/gt}$ retinas. (**A**) Detection of apoptotic cells (TUNEL, green) and corresponding quantification (right) in young, middle-aged, and old *Ambra1*$^{+/+}$ and *Ambra1*$^{+/gt}$ mice (n = 3–9). (**B**) Immunostaining of red/green cones (RG opsin, cyan; left) and corresponding quantification (right) in temporal to nasal regions in old *Ambra1*$^{+/+}$ and *Ambra1*$^{+/gt}$ mice *(*n = 5). **(C)** Immunostaining of microglial cells (IBA1, magenta) in retinal flat mounts (left) from old *Ambra1*$^{+/+}$ and *Ambra1*$^{+/gt}$ littermates. Two distinct microglial cell morphologies (ramified and ameboid) were observed (right) (n = 5–6 per group). Arrows indicate ameboid microglial cells. (**D**) Representative images showing immunostaining of gliosis (GFAP, magenta) and Müller cells (GS, green) in young, middle-aged, and old *Ambra1*$^{+/+}$ (left panels) and *Ambra1*$^{+/gt}$ mice (right panels). Nuclei are counterstained with DAPI (grey). Data are presented as the mean ± SEM. $^{*}p < 0.05$: two-way ANOVA followed by Fisher's LSD *post hoc* for genotype (**B**). Scale bars: 50 µm (**A, C, D**); 25 µm (**B**).

### Supplementary Figure S4

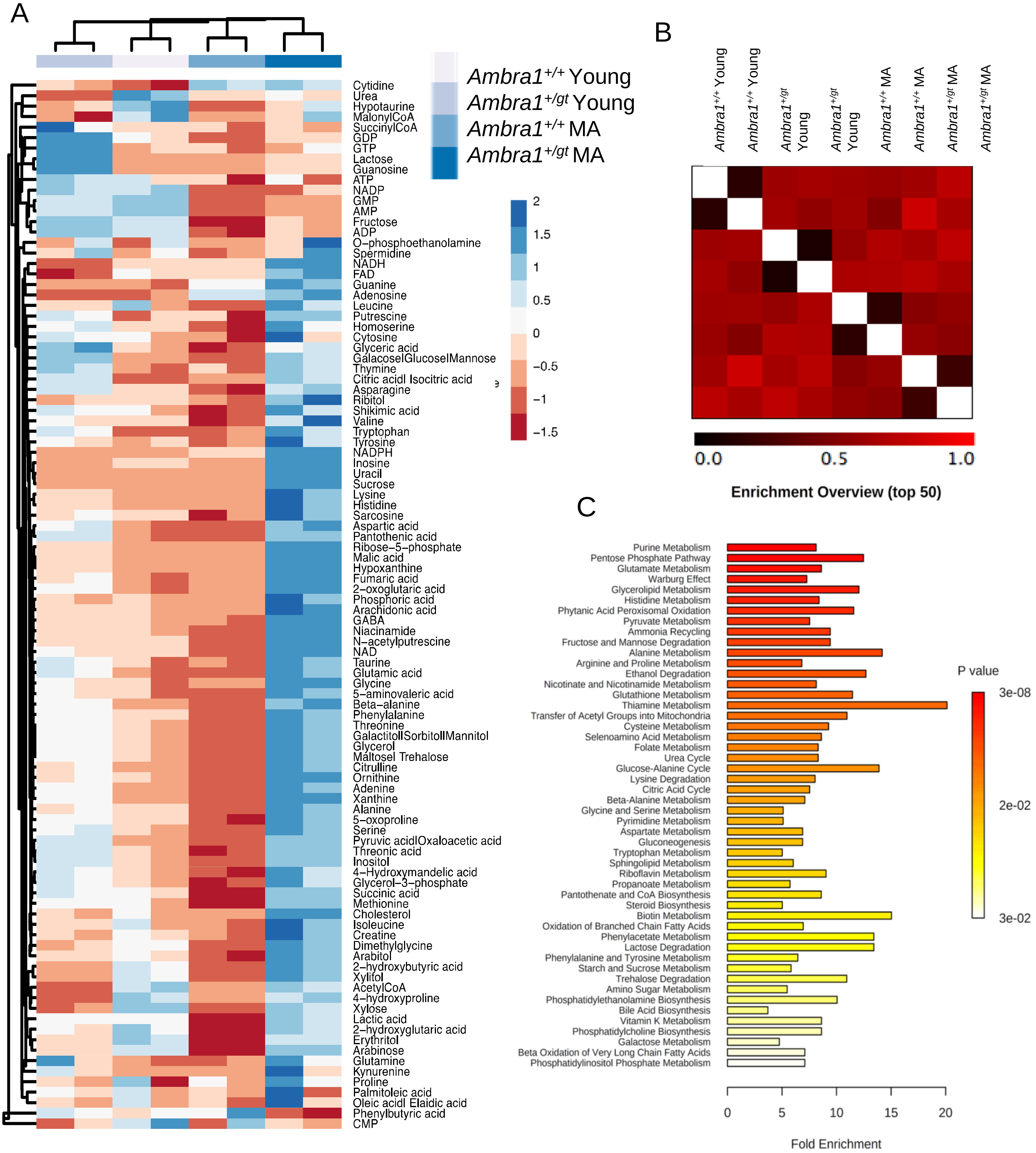


Supplementary Figure S4. Detailed metabolomic analysis of *Ambra1*$^{+/gt}$ retinas. (**A**) Heat map and hierarchical clustering (HCL) analysis of all metabolites from young and middle-aged (MA) *Ambra1*$^{+/+}$ and *Ambra1*$^{+/gt}$ retinas (n = 2 per group). (**B**) Pairwise Pearson´s correlation of biological replicates from a metabolomic study of whole retina extracts from young and middle-aged (MA) *Ambra1*$^{+/+}$ and *Ambra1*$^{+/gt}$ mice (n = 2 per group). (**C**) Pathway enrichment analysis of metabolites for which significant differences were observed between *Ambra1*$^{+/+}$ and *Ambra1*$^{+/gt}$ duplicates (n = 2 per group). Statistical significance (Skillings-Mack test) was set at $p < 0.05$.

Supplementary Figure S5

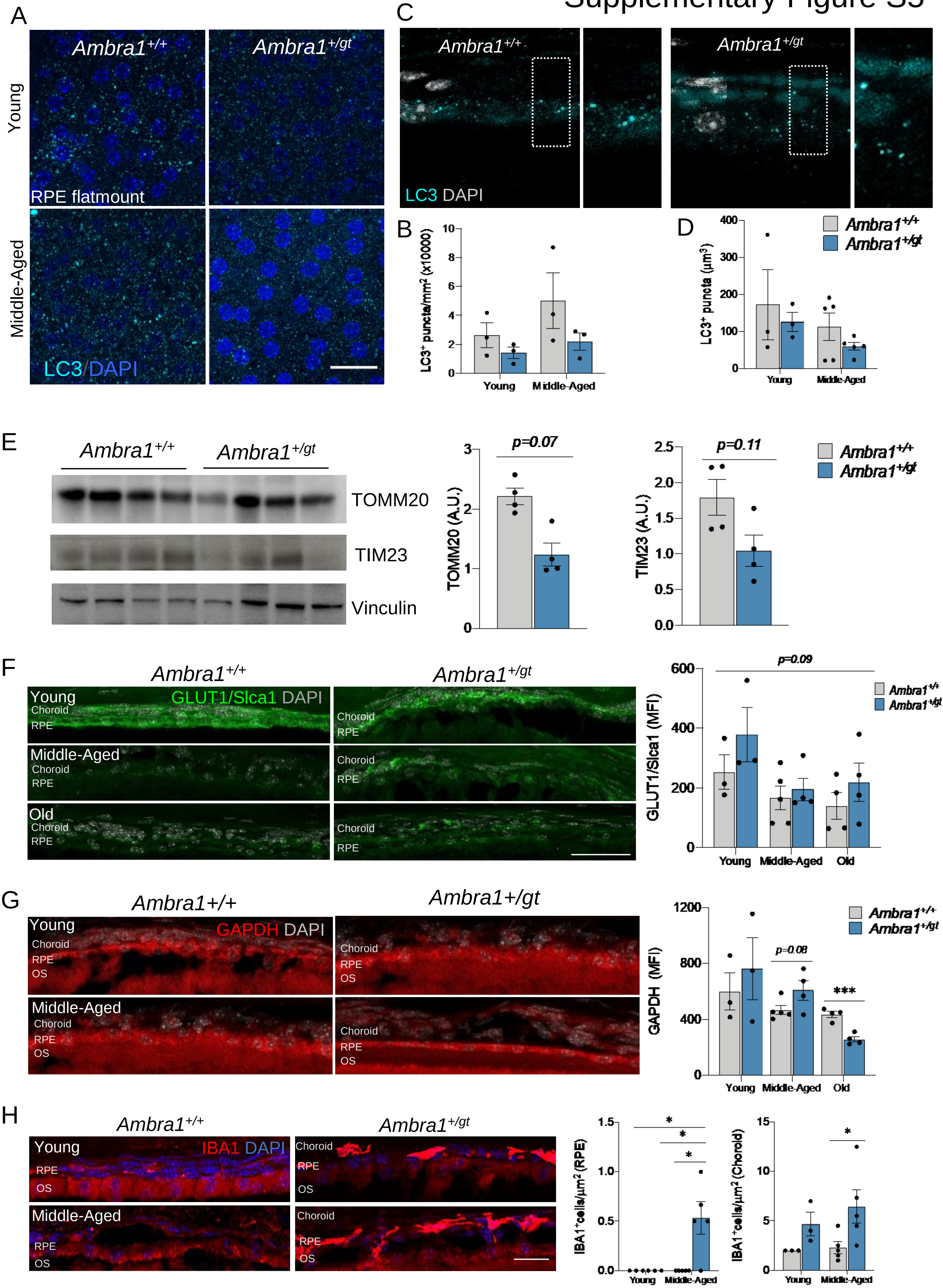

Supplementary Figure S5: Middle-aged *Ambra1*$^{+/gt}$ mice show alterations in autophagy and metabolism in the RPE. (**A, B**) RPE flat mounts from *Ambra1*$^{+/+}$ and *Ambra1*$^{+/gt}$ littermates immunostained for LC3 and corresponding quantification (n = 3 per group). (**B**) Quantification of LC3+ puncta in **A**. (**C, D**) Immunostaining for LC3 in RPE cryosections from *Ambra1*$^{+/+}$ and *Ambra1*$^{+/gt}$ littermates and corresponding quantification of LC3+ puncta in young and middle-aged animals (n = 5 per group). (**E**) Levels of TOMM20 and TIM23 proteins in the RPE were evaluated by Western blot in middle-aged *Ambra1*$^{+/+}$ and *Ambra1*$^{+/gt}$ mice. (**F**) Immunostaining of GLUT1 in RPE cryosections in young, middle-aged, and old *Ambra1*$^{+/+}$ and *Ambra1*$^{+/gt}$ littermates. Corresponding quantification of GLUT1 mean fluorescence intensity (n = 3–5 per group). (**G**) Immunostaining of GAPDH in RPE cryosections from young and middle-aged *Ambra1*$^{+/+}$ and *Ambra1*$^{+/gt}$ littermates. Corresponding quantification of GAPDH in young, middle-aged, and old *Ambra1*$^{+/+}$ and *Ambra1*$^{+/gt}$ littermates (n = 3–5 per group). (**H**) Immunostaining of microglial cells (IBA1, red) (left) in the RPE and choroid of young and middle-aged *Ambra1*$^{+/+}$ and *Ambra1*$^{+/gt}$ mice (n = 3–5 per group). Corresponding quantification is shown on the right. Nuclei were counterstained with DAPI (blue or grey). Data are presented as the mean ± SEM. *$p < 0.05$; two-way ANOVA followed by Fisher's LSD *post hoc* test for genotype (**B, D**); two-tailed Student's *t*-test (E, TIM23); or Mann Whitney *U*-test (E, TOMM20). Scale bars: 50 µm € and 25 µm (**A**, **F**, **G** and **H**).

### Supplementary Figure S6

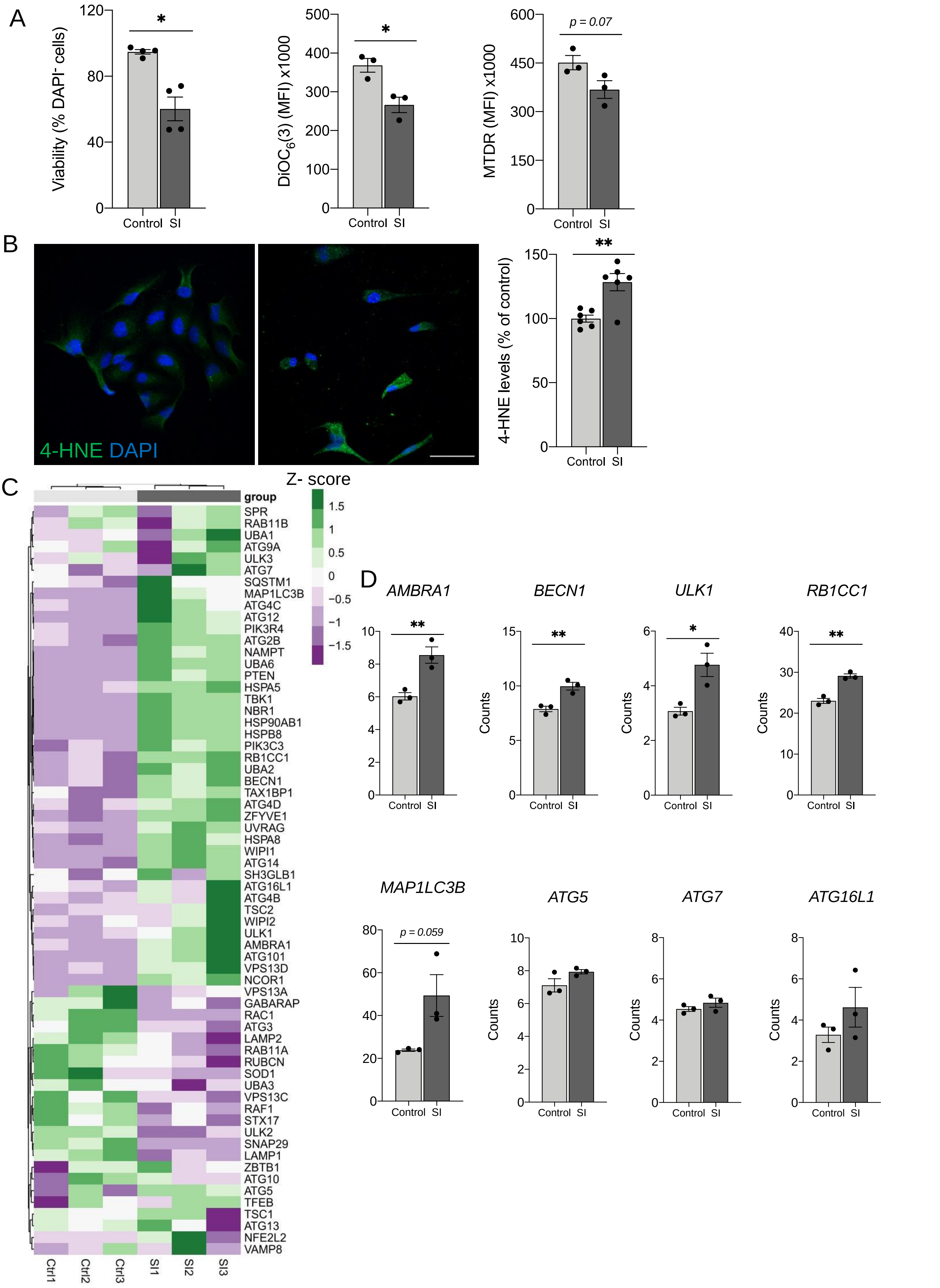

Supplementary Figure S6: Sodium iodate (SI) induces cell death and transcription of autophagy machinery genes in ARPE19 cells. (**A**) ARPE-19 cells were treated with 20 mM SI for 24 h and then analyzed by flow cytometry to assess viability (DAPI exclusion assay), ΔΨm ($DiOC_6(3)$), and mitochondrial mass (MitoTracker Deep Red, MTDR). n = 3–4 independent experiments with 2 biological replicates. (**B**) ARPE-19 cells were treated with 20 mM SI for 24 h and lipid peroxidation was assessed by immunofluorescence for 4-HNE (green). Nuclei were counterstained with DAPI (blue). n = 6 biological replicates from 2 independent experiments. (**C, D**) RNA-seq dataset (GSE142591) from ARPE19 cells treated with 20 mM SI for 24 h. (**C**) A manually-curated autophagy gene list was used to generate a heatmap of unsupervised hierarchical clustered samples and genes (n = 3). (**D**) Gene expression based on normalized counts of selected autophagy genes (*AMBRA1, BECN1, ULK1, RB1CC1, MAP1LC3B, ATG5, ATG7, ATG16L1*) from the same dataset (n = 3). Data are presented as the mean ± SEM. *p<0.05, **p<0.01: two-tailed Student's *t*-test (**A**, **B**, **D**). Scale bar: 50 μM.

### Supplementary Figure S7

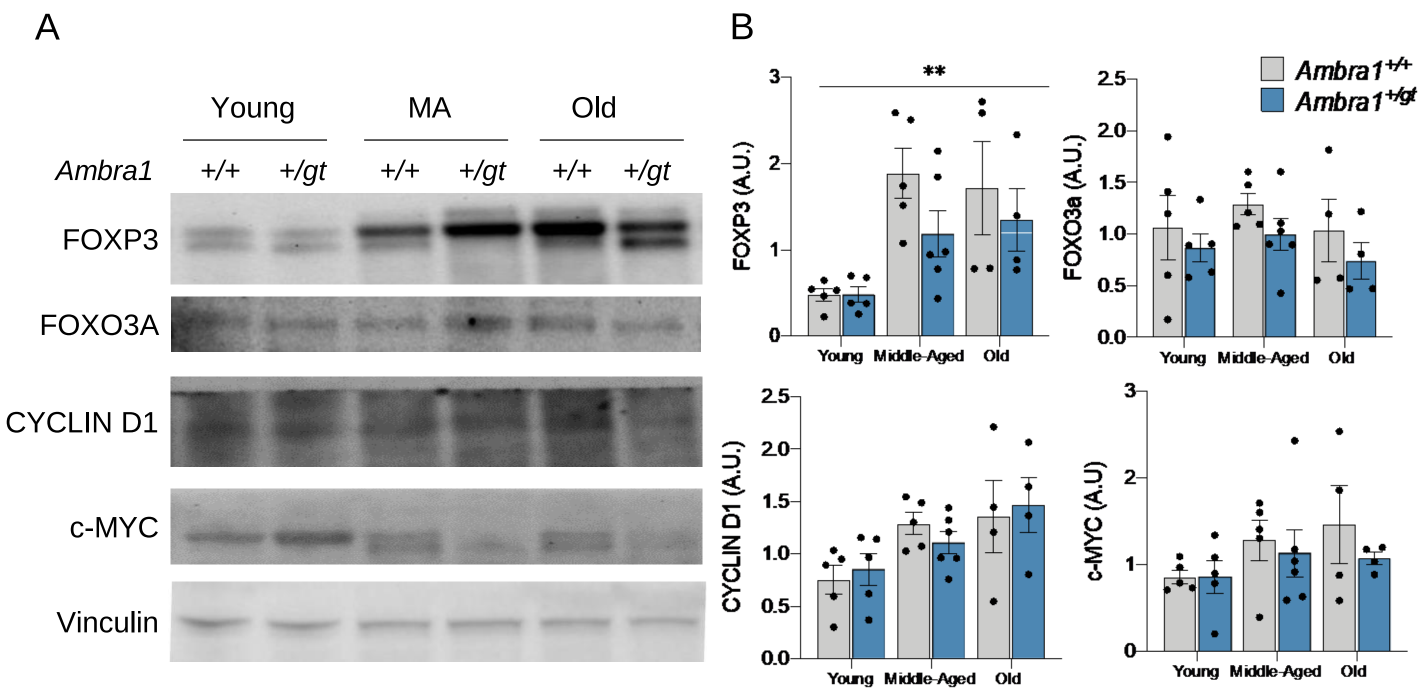


Supplementary Figure S7: Non-autophagic functions of Ambra1 are unchanged in the RPE of *Ambra1*$^{+/gt}$ mice. (**A**) Protein levels of FOXP3, FOXO3A, CYCLIN D1 and c-MYC were evaluated by Western blot in the RPE in young, middle-aged, and old *Ambra1*$^{+/+}$ and *Ambra1*$^{+/gt}$ mice. (**B**) Quantification of Western blots shown in A, expressed as protein level relative to that of the loading control (vinculin) (n = 4–5 per age/genotype). Statistical analysis was performed by 2-way ANOVA followed by Fisher's LSD *post hoc* test. *p<0.05.

### Supplementary Figure S8

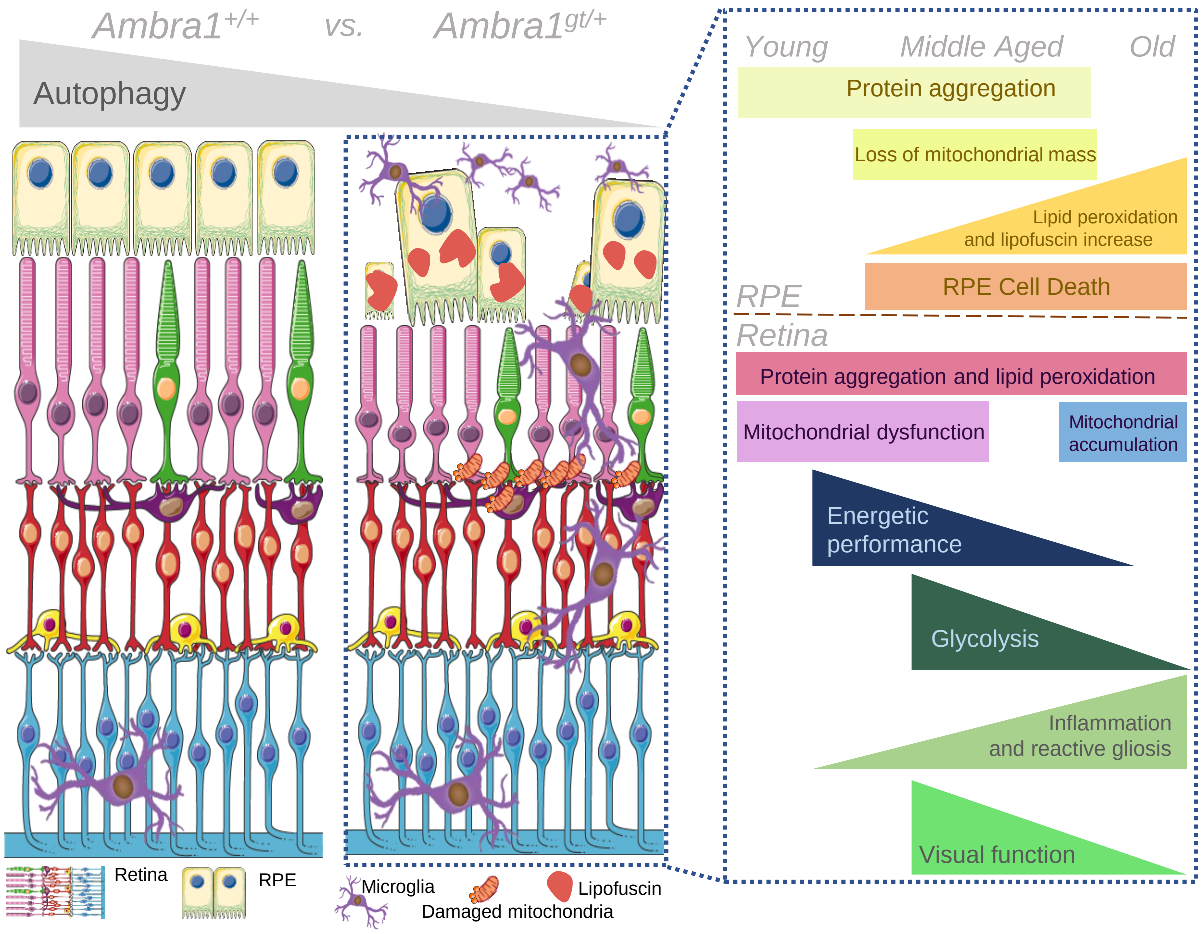


Supplementary Figure S8: Graphical abstract. *Ambra1*$^{+/gt}$ mice display diminished autophagy activity in both the retina and RPE. This decrease in autophagy is accelerated relative to that which occurs during physiological aging. This age-associated autophagy deficiency results in premature degeneration of the RPE and retina and loss of visual function. Cellular and functional degeneration the RPE-retina is associated with protein accumulation, oxidative stress, and inflammation. Cellular damage disrupts the metabolic balance of the RPE-retina, leading to reduced mitochondrial mass in degenerating RPE cells and mitochondrial dysfunction and reduced glycolysis in the retina. Graphical abstract images templates were obtained from Servier Medical Art (https://smart.servier.com/).
